# A 3’ UTR-derived small RNA connecting nitrogen and carbon metabolism in enteric bacteria

**DOI:** 10.1101/2022.04.01.486790

**Authors:** Lauren R. Walling, Andrew B. Kouse, Svetlana A. Shabalina, Hongen Zhang, Gisela Storz

**Author notes:** The authors wish it to be known that, in their opinion, the first two authors should be regarded as Joint First Authors. Present address: Andrew Kouse, Catalent Pharma Solutions, Baltimore, MD 21201, USA.

## Abstract

Increasing numbers of small, regulatory RNAs (sRNAs) corresponding to 3’ untranslated regions (UTR) are being discovered in bacteria. One such sRNA, denoted GlnZ, corresponds to the 3’ UTR of the *Escherichia coli glnA* mRNA encoding glutamine synthetase. Several forms of GlnZ, processed from the *glnA* mRNA, are detected in cells growing with limiting ammonium. GlnZ levels are regulated transcriptionally by the NtrC transcription factor and post-transcriptionally by RNase III. Consistent with the expression, *E. coli* cells lacking *glnZ* show delayed outgrowth from nitrogen starvation compared to wild type cells. Transcriptome-wide RNA-RNA interactome datasets indicated that GlnZ binds to multiple target RNAs. Immunoblot and assays of fusions confirmed GlnZ-mediated repression of *glnP* and *sucA*, encoding proteins that contribute to glutamine transport and the citric acid cycle, respectively. Although the overall sequences of GlnZ from *E. coli* K-12, Enterohemorrhagic *E. coli* and *Salmonella enterica* have significant differences due to various sequence insertions, all forms of the sRNA were able to regulate the two targets characterized. Together our data show that GlnZ promotes survival of *E. coli* under low nitrogen conditions by modulating genes that affect carbon and nitrogen flux.

## INTRODUCTION

Nitrogen is central to the metabolic needs of all organisms and is required for most cellular components. Consequently, nitrogen starvation reduces growth (reviewed in (1, 2)). Maintenance of intracellular nitrogen levels to avoid starvation while preventing the wasteful production of transporters or biosynthetic machinery requires proper regulation (reviewed in (1,3,4)).

*E. coli* satisfies its nitrogen requirements through the import of nitrogen sources from its surrounding environments. The preferred nitrogen source ammonium is transported into the cytoplasm and converted into glutamine, one of the major nitrogen sinks in *E. coli* (reviewed in (5)). Ammonium first reacts with *α*-ketoglutarate to give glutamate, in a reaction catalyzed by the GdhA glutamate dehydrogenase. Glutamate can then combine with another molecule of ammonium to give glutamine, in a reaction catalyzed by the GlnA glutamine synthetase. Glutamine also is transported into the cytoplasm by the GlnPQ ABC transporter (reviewed in (3)). When ammonium and glutamine are not available in the extracellular environment, *E. coli* can upregulate the transport and catabolism of nucleotides, amino acids or other metabolites to obtain nitrogen (reviewed in (6, 7)). Given the importance of GdhA and GlnA and their connections to various aspects of bacterial metabolism, the expression and activities of these enzymes is regulated at multiple levels (reviewed in (3)).

Transcription of *gdhA*, *glnA* and other genes encoding transporters of alternative nitrogen sources, catabolic machinery and biosynthetic machinery used by *E. coli* to satisfy its nitrogen requirements is regulated by the NtrB (GlnL/NRII)-NtrC (GlnG/NRI) two-component response system together with σ^54^ (RpoN) (reviewed in (6–8)). When cells encounter nitrogen limitation, the NtrBC response is activated and promotes the σ^54^-dependent activation of these genes (reviewed in (3)).

In addition to being controlled by transcription factors, many responses to changes in environmental conditions are modulated by small regulatory RNAs (sRNAs). In *E. coli*, most sRNAs act by limited base pairing with target mRNAs in a manner that is facilitated by the Hfq RNA chaperone protein (reviewed in (9)). Base pairing with the target mRNAs can lead to decreased or increased translation and/or can have effects on target RNA transcription and/or stability (reviewed in (10)). sRNAs initially were discovered as independent transcription units encoded in intergenic regions. However, more recently, increasing numbers of sRNAs have been found to correspond to the 3’ untranslated regions (UTRs) of protein coding genes, transcribed from promoters internal to the coding sequences or processed from the mRNAs or both (reviewed in (11, 12)). While a number of *E. coli* sRNAs, such as the Spot 42 and GlmZ sRNAs, regulate carbon metabolism (reviewed in (13)), only few sRNAs have been found to affect nitrogen metabolism. SdsN, which is induced in stationary phase, and NarS, which is induced by anaerobic shock, protect against the deleterious nitrogen compounds nitrofurans and nitrite, respectively (14, 15).

Here we describe the characterization of another sRNA that impacts nitrogen as well as carbon metabolism and is derived from the 3’ UTR of the *glnA* gene encoding glutamine synthetase. This sRNA, originally denoted *glnA* 3’ UTR, but renamed GlnZ, was first identified in a cloning-based screen that identified multiple short 5’ and 3’ UTR-derived transcripts in *E. coli* (16). GlnZ overlaps the *glnA* ORF and extends into the intergenic region between *glnA* and *glnLG* (encoding NtrB and NtrC, respectively). We examined GlnZ expression, phenotypes associated with a *glnZ* deletion, effects of the sRNA on predicted target genes and its conservation. These experiments showed GlnZ represses the synthesis of a glutamine transporter and central metabolic enzymes and promotes survival during nitrogen starvation. Interestingly, although there are significant length and sequence differences in the 3’ UTRs of *glnA* of *E. coli* K12, Enterohemorrhagic *E. coli* (EHEC) O157:H7 and *Salmonella enterica* LT2, sRNAs containing highly similar seed and terminator sequences, are generated from all these regions in low nitrogen and likely repress overlapping targets.

## MATERIALS AND METHODS

### Bacterial strains and plasmids

Bacterial strains, plasmids and oligonucleotides used in this study are listed in Supplementary Tables S1, S2 and S3, respectively. All MG1655 WT strains are *crl^+^* (GSO982) except where noted. Strains constructed in this study were created by recombineering following published procedures (17, 18). Briefly, deletions were constructed by replacing the sequence of interest with a kanamycin cassette using the *λ* red recombinase system in the NM400 *E. coli* strain. The deletions were then moved into the MG1655 *E. coli* strain by P1 transduction. The chromosomal *sucA-M1-SPA* mutant was constructed by replacing the *sucA* sequence with a *cat-sacB* cassette using the *λ* red recombinase system in the NM500 *E. coli* strain. The same recombineering system was then used to replace the *cat-sacB* cassette with either *sucA-SPA*::*kan* or *sucA-M1- SPA*::*kan*. The alleles were then moved into the MG1655 *E. coli* strain by P1 transduction.

Plasmids were constructed by amplifying the sequence of interest using the MG1655 genomic DNA as template and inserting it into the desired vector using either the Gibson Assembly Cloning Kit (New England Biolabs) or by digestion with the indicated restriction enzymes followed by ligation using T4 DNA ligase (New England Biolabs). All strain mutations and plasmid inserts were confirmed by sequencing.

### Bacterial growth conditions

Unless specified otherwise, cultures were grown aerobically overnight at 37°C in LB (10 g tryptone, 5 g yeast extract, 5 g NaCl/l). Overnight cultures were subcultured 1:100 in Gutnick medium (4.6 g KH_2_PO_4_, 13.5 g K_2_HPO_4_, 1 g K_2_SO_4_ and 0.1 g MgSO_4_-7H_2_O/l) supplemented 1:1000 with Ho-LE (2.86 g H_3_BO_3_, 1.80 g MnCl_2_•4H_2_O, 1.36 g FeSO_4_, 1.77 g C_4_H_4_O_6_Na_2_, 26.90 mg CuCl_2_.2H_2_O, 20.80 mg ZnCl_2_, 40.40 mg CoCl_2_•6H_2_O, 25.20 mg Na_2_MoO_4_•2H_2_O/l) (19) with the indicated carbon and nitrogen sources and incubated aerobically at 37°C. Antibiotics were used at the following final concentrations: ampicillin at 100 μg/ml, kanamycin at 30 μg/ml, chloramphenicol at 25 μg/ml. Arabinose was supplemented as 0.2% of total culture volume and rifampicin was added at a final concentration of 50 μg/ml.

### RNA extraction

RNA was extracted from *E. coli* cells using the TRIzol reagent (Thermo Fisher Scientific) as previously described (14). Briefly, 10 OD_600_ of cells were collected and then resuspended in 1 ml of TRIzol reagent. Samples were incubated at room temperature for 5 min, 0.2 ml of chloroform was added, and samples were vortexed. Samples were collected at maximum speed for 15 min at 4°C, and 0.6 ml of the aqueous layer was transferred to a fresh Eppendorf tube with equal volume of isopropanol. Following a 10-min room temperature incubation, samples were centrifuged at maximum speed for 15 min at 4°C to collect the RNA pellet, which was washed wit h 1 ml of 70% ethanol. RNA pellets were air dried and dissolved in DEPC-treated water.

Total RNA concentrations were determined by OD_260_.

### Northern blot analysis

Northern blot analysis was performed as previously described (14). Briefly, 10 μg of total RNA was loaded into each well of an 8% polyacrylamide–7M urea gel (USB Corporation) run at 300 V for 1.5 h in 1x TBE. The gel was transferred overnight to Zeta-Probe membrane (Bio-Rad) at 12 V at 4°C. Following transfer, membranes were UV crosslinked and pre-hybridized in ULTRAhyb Oligo Hybridization Buffer (Invitrogen) for 1 h for at 45°C. Oligonucleotides complimentary to the target were end-labeled with γ-^32^P-ATP by T4 polynucleotide kinase (New England Biolabs). Labeled oligonucleotides were added to the membrane and incubated overnight at 45°C. Subsequently, membranes were rinsed 2X with 2× SSC + 0.1% SDS, 1X with 0.2× SSC + 0.1% SDS, washed with 0.2× SSC + 0.1% SDS at 45°C for 25 min and then rinsed 1X with 0.2× SSC + 0.1% SDS. Membranes were exposed to KODAK Biomax X-ray film at - 80°C.

### Immunoblot analysis

Samples were grown under the conditions indicated and the same number of cells were resuspended in 2× protein loading buffer. Proteins were separated on a 15% denaturing SDS-PAGE gel (run at 200V for 30 min in 1× Tris Glycine SDS Running Buffer). Samples were then transferred to a 0.2 μM pore size nitrocellulose membrane (Novex) using 1× transfer buffer with 30% methanol. Following transfer, the membrane was blocked in 5% milk in TBS with 0.1% Tween-20 for 1 h at room temperature. For the detection of SucA-SPA, the blots were incubated 1 h at room temperature with a 1,2000 dilution of anti-FLAG(M2)-HRP (Sigma) in TBST with 5% milk. For the detection of glutamine synthetase, we used rabbit antiserum (a kind gift of Rodney Levine, National Heart, Lung, and Blood Institute) raised against recombinant *E*. *coli* glutamine synthetase. The blot was incubated overnight at 4°C with a 1:5,000 dilution of the antiserum in TBST with 5% milk, followed by a 1 h incubation at room temperature with a 1:5,000 dilution of secondary anti-rabbit-HRP antibody (Invitrogen) in TBST. For both the anti-FLAG and anti-GlnA blots, the membranes were then incubated for 1 min with SuperSignal West Pico (Thermo Fisher Scientific) detection reagent and imaged.

### Nitrogen starvation growth assay

Cultures grown overnight in Gutnick medium with Ho-LE, 15 mM ammonium and 0.4% glucose were subcultured 1:100 in Gutnick medium with Ho-LE, 3 mM ammonium and 0.4% glucose.

These cultures were grown at 37°C to nitrogen starvation (6 h) and then incubated for an additional 48 h. Following this period of starvation, cells were subcultured to an OD_600_ of 0.05 in Gutnick medium with Ho-LE, 3 mM ammonium and 0.4% glucose. Growth was assayed by measuring OD_600_ every 10 min on a CLARIOstar plate reader.

### β-galactosidase assay

Cultures carrying the *lacZ* fusion of interest were cultured in Gutnick medium with Ho-LE, 15 mM ammonium, 0.4% glucose and 0.2% arabinose. After, cultures were grown for the indicated times, OD_600_ was measured, and 100 μl of cells were transferred to 700 μl of Z-buffer, 30 μl of chloroform and 15 μl of 0.1% SDS. After vortexing and incubation at 28°C for 15 min, 100 μl of 8 mg/ml ONPG in Z-buffer was added to each sample. Once samples turned slightly yellow, the reactions were stopped by the addition of 500 μl of 1M Na_2_CO_3_. *β*-galactosidase activity was calculated based on OD_420_ and OD_550_ measurements using the established formula (20).

### RNA structure probing

GlnZ_194_ RNA was *in vitro* transcribed using a MEGAShortscript T7 transcription kit (Invitrogen). After the sample was treated with DNase I for 15 min at 37°C, ammonium acetate stop solution was added, and the RNA was extracted with phenol:chloroform:isoamylalcohol, followed by precipitation in 100% ethanol with sodium acetate and Glycoblue (Invitrogen). The RNA pellet was washed with 70% ethanol, dried, and resuspended in 30 µl DEPC water. The quality and concentration of RNA was checked on a 4200 TapeStation (Agilent Technologies). The RNA (50 pmol) was dephosphorylated by CIP treatment, re-purified as described above and then radiolabeled with *γ*[^32^P]-ATP and T4 PNK. Unincorporated *γ*[^32^P]-ATP was removed with a G-50 spin column, and the radiolabeled RNA was resolved on an 8% polyacrylamide gel. The RNA was then eluted from the gel by an overnight 4°C incubation in RNA elution buffer (0.1 M sodium acetate, 0.1% SDS, 10 mM EDTA). RNA was purified from the supernatant, as described above.

To perform structure probing, RNA (0.2 pmol) was denatured at 95°C for 1 min, followed by 5 min on ice. After the addition of yeast RNA and RNA structure buffer (Invitrogen), samples were renatured by incubation at 37°C for 1 h. Next samples were treated with either 1 µl of RNase III or RNase T1 (0.01 U/µl) and incubated at 37°C for 0, 3, 5, or 10 min. Reactions were stopped by the addition of 20 µl of Inactivation/Precipitation Buffer (Invitrogen). For the OH ladder, 0.4 pmol of labeled RNA was combined with 9 µl of alkaline hydrolysis buffer and incubated 5 min at 90°C. For the T1 ladder, 0.4 pmol of labeled RNA was denatured and then incubated with 0.1 U RNase T1 for 5 min at 37°C. RNA was resolved on an 8% acylamine/8 M urea gel (70 Watts). The gel was dried using a ThermoSavant Stacked Gel Dryer SGD300 and exposed to film.

### MAPS analysis

Strains containing either pNM-MS2 vector or pNM-MS2-GlnZ_194_ were grown in 100 ml of Gutnick medium supplemented with HoLE, 0.4% glucose, and 15 mM glutamine to stationary phase (∼8 h), then treated with 1 mM IPTG for 1 h. Cells were pelleted and snap frozen in liquid nitrogen, then resuspended in 4 ml of ice-cold Buffer A (150 mM KCl, 20 mM Tris-HCl pH 8.0, 1 mM MgCl_2_, 1 mM DTT) and sonicated to lyse the cells. Cell debris was removed by centrifugation (4,000 rpm for 10 min), and the supernatant was used for affinity chromatography, except 150 µl for lysate-control RNA. Poly-Prep chromatography columns were loaded with 300 µl of amylose resin (New England Biolabs). After columns were washed 3X with 10 ml of cold Buffer A, MS2-MBP (3,600 pmol diluted in 6 ml of cold Buffer A) was added. Cell lysates were loaded on the columns, and the columns were washed 3X with 10 ml of cold Buffer A, followed by elution in 1 ml of cold Buffer E (250 mM KCl, 20 mM Tris-HCl pH 8.0, 12 mM maltose, 0.1% Triton, 1 mM MgCl_2_, 1 mM DTT). RNA was extracted by adding 1 ml Trizol reagent as described above.

After quantification using the 4200 TapeStation, RNA was treated with recombinant RNase inhibitor (40 U), followed by fragmentation and de-phosphorylation by the addition of 4 µl of 10X FastAP buffer and incubation for 3 min at 94°C. Samples were placed on ice and 20 µl of DNase-phosphatase reaction mixture (1 U/µl recombinant RNase inhibitor, 0.2 U/µl TURBO DNase, 0.25 U/µl FastAP) was added. After a 30-min incubation at 37°C, the RNA was purified using the RNA Clean & Concentrator-5 kit (Zymo Research). The 3’ barcoded adaptors were ligated onto the RNA by mixing 5 µl of the dephosphorylated RNA with 100 µM of the barcoded adaptor and heating for 2 min at 70°C before transfer to ice. Next, 14 µl of ligation mix (1X T4 RNA ligase buffer, 9% DMSO, 1 mM ATP, 20% PEG 8000, 0.6 U/µl recombinant RNase inhibitor, and 2.55 U/µl T4 RNA ligase 1) was added to all samples, which were incubated at 22°C for 2 h. The ligation reaction was stopped by the addition of 60 µl of RLT buffer (Qiagen). All samples were then pooled and cleaned using the RNA Clean & Concentrator-5 kit (Zymo Research).

Ribosomal RNA was depleted using the RiboCop kit (Lexogen) according to the manufacturer’s instructions. First-strand cDNA synthesis was performed using the SuperScript III First-Strand Synthesis system (Invitrogen). RNA was then degraded by adding 2.5 µl of 1N NaOH and incubating for 12 min at 70°C, followed by the addition of 5 µl of 0.5 M acetic acid. A second adaptor was ligated at the cDNA 3’ end using T4 RNA ligase (2.25 U/µl) and an overnight incubation at 22°C, whereupon the sample was amplified by PCR. Between each of the steps the samples were purified using AMPure XP Beads (Beckman Coulter). The final PCR product was analyzed using the Qubit dsDNA HS Assay Kit (ThermoFisher) according to the manufacturer’s instructions and using a TapeStation (Agilent) and High-Sensitivity D1000 ScreenTape (Agilent).

RNA-seq libraries were sequenced on the Illumina HiSeq 2500 platform at Molecular Genomics Core, *Eunice Kennedy Shriver* National Institute of Child Health and Human Development, National Institutes of Health, Bethesda, USA. The raw fastq records were demultiplexed with python script index_splitter.py (https://github.com/asafpr/RNAseq_scripts/blob/master/index_splitter.py) and trimmed with cutadpt software (version 3.0) to remove adapter and internal barcode sequences. The trimmed fastq reads were mapped to *E. coli* genome (ecoli-k12-MG1655-NC_000913-3) with Python RILSeq package (version 6.0, https://github.com/asafpr/RILseq). Reads counts were obtained with featureCounts tool of Subread software (version 2.0.3) and customized GTF file which annotates all of gene/exon, 5’ UTR, 3’ UTR, and intergenic regions. Differential expression analyses were conducted with R DESeq2 package with default normalization procedures and apeglm algorithm for log2 fold change shrinkage (21, 22).

### Fluorescence assays

Cultures grown overnight in LB were subcultured 1:100 into fresh LB with ampicillin, arabinose, and IPTG and then grown at 37°C for 3 h to an OD_600_ ∼1.0. At this time, 1 ml of culture was removed and pelleted by centrifugation at maximum speed for 1 min. The pellet was resuspended in 300 µl of PBS, and 100 µl of each sample was placed into a black 96-well plate to measure mCherry activity (excitation at 555 nm, emission at 635 nm) or YFP activity (excitation at 497 nm, emission at 540 nm) on a PerkinElmer EnVision 2105 plate reader (mCherry) or a CLARIOstar plate reader (YFP). For normalization, OD_600_ was measured for 100 µl of each sample in a clear 96-well plate. An average of three biological replicates is reported.

### Primer extension analysis

Cultures grown in Gutnick medium with HoLE, 0.4% glucose, 15 mM ammonium, ampicillin, and 1 mM IPTG to late logarithmic phase (OD_600_ ∼1.5) were collected, and RNA was isolated as described above. Primer LW055 was radiolabeled as described above, and was annealed to RNA by incubating with 5 µg RNA at 80°C, slowly decreasing to 42°C. Each sample was then incubated with 0.5 µl AMV Reverse Transcriptase (Life Sciences Advanced Technologies Inc) and 2.5 µM of each dNTP for 1 h at 42°C. The sequencing ladder was generated using a Sequenase kit (Thermo Fisher) with aceE DNA amplified by PCR as the template, following the protocol provided. RNA was resolved on an 8% acylamide/8 M urea gel (70 Watts). The gel was dried using a ThermoSavant Stacked Gel Dryer SGD300 and exposed to film.

### Evolutionary analysis and comparison of GlnZ architecture

The sequences of 3’ UTRs of the conserved glutamine synthetase (*glnA*) gene of various bacterial species were isolated based on the genome annotation provided in GenBank (https://www.ncbi.nlm.nih.gov /search/all/?term=GenBank). To identify orthologous sequences, we applied Blast (23), where the search parameters were adjusted to search for a short input sequence, such as ∼20-25 nt of the seed or terminator regions of GlnZ.

MUSCLE alignments (24) and edited to take into account the results of pairwise comparisons, which was done using the OWEN program (25). Three GlnZ classes were identified based on the search software (26, 27). GlnZ sequences were computationally folded using the Afold program (28) to predict conserved stable and relaxed structures. This program also was applied to estimate the duplex free energy between GlnZ seed sequences and their targets in all three classes. The predicted minimum free energies for optimal and suboptimal structures were calculated using an implementation of the dynamic programming algorithm (29), which employs nearest neighbor parameters to estimate free energy.

Tamura-Nei model (31). Initial tree(s) for the heuristic search were obtained automatically by applying Neighbor-Join and BioNJ algorithms to a matrix of pairwise distances estimated using the Maximum Composite Likelihood (MCL) approach, and then selecting the topology with superior log likelihood value. A discrete Gamma distribution was used to model evolutionary rate differences among sites (5 categories (+*G*, parameter = 0.9684)). The rate variation model allowed for some sites to be evolutionarily invariable ([+*I*], 37.79% sites).

## RESULTS

### GlnZ levels increase upon nitrogen starvation

The GlnZ sRNA was first identified in a screen for 3’ UTR-derived sRNAs in *E. coli* (16). A genome-wide analysis of 5’ ends by differential RNA-seq (32) indicated a promoter internal to the *glnA* coding sequence with two additional processed 5’ ends, likely generated by RNase cleavage. Given the 3’ end detected past a predicted intrinsic terminator between *glnA* and *glnL* in Term-seq data (33), these 5’ ends should give RNA products of 213 nt (from the predicted promoter) and 194 and 174 nt (from the two processed ends) (Figure 1A).

**Figure 1.**
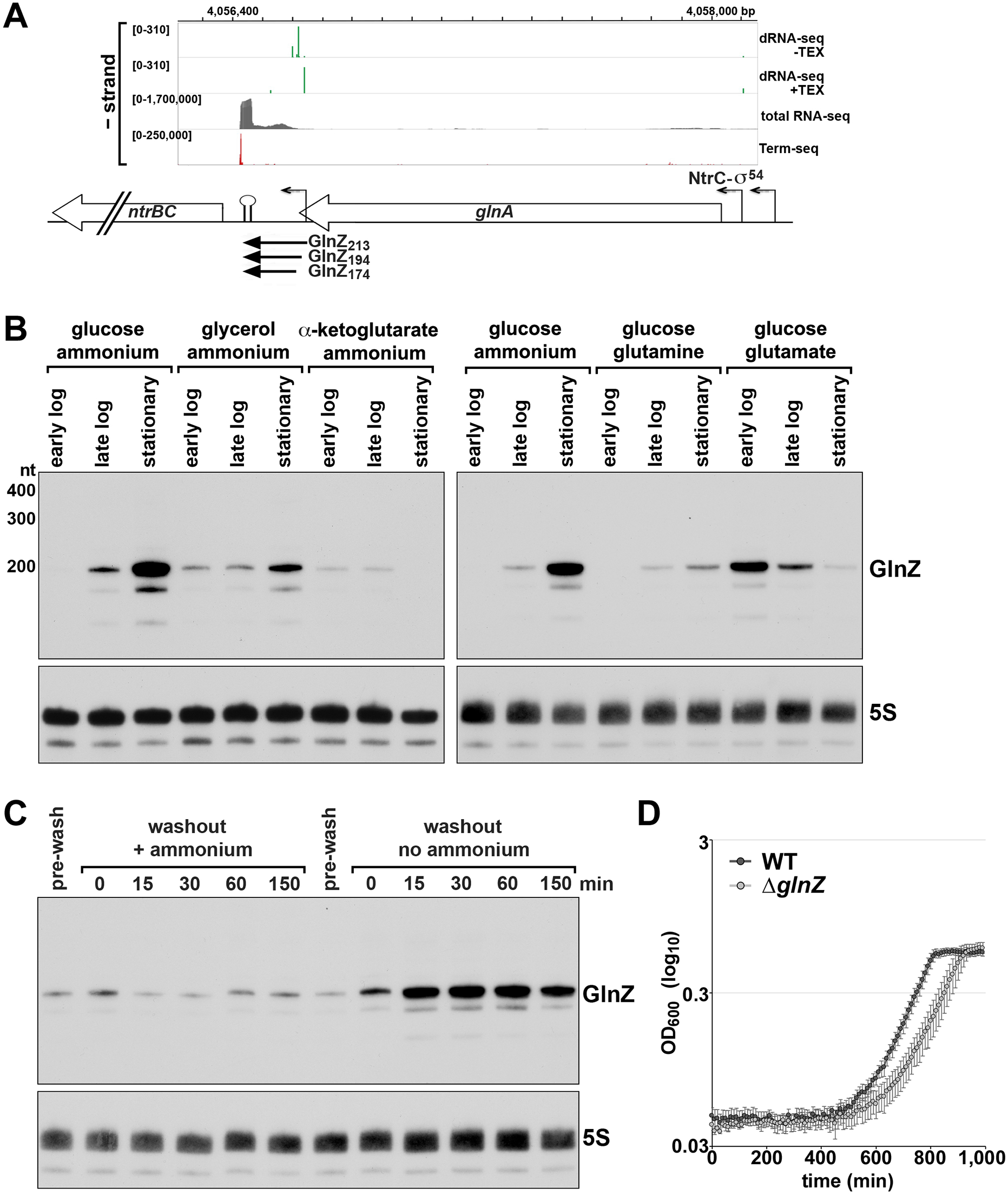
GlnZ expression across growth in different carbon and nitrogen sources. (**A**) Organization of the genomic region encoding GlnZ. dRNA-seq data with or without Terminator Exonuclease (TEX) treatment (32) is depicted in green. Total RNA-seq and Term-seq data (33) are depicted in grey and red, respectively. Only GlnZ_213_ includes the glnA stop codon. (B) Northern blot analysis of GlnZ levels with WT MG1655 cultured to early logarithmic, late logarithmic or stationary phases in Gutnick medium with 0.4% glucose, glycerol or *α*-ketoglutarate as the sole carbon source and 15 mM ammonium as the nitrogen source or Gutnick medium with 0.4% glucose as the sole carbon source and 15 mM ammonium, glutamine or glutamate as the nitrogen source. (**C**) Northern blot analysis of GlnZ levels with WT MG1655 cultured to mid logarithmic phase in Gutnick medium with 0.4% glucose and 15 mM ammonium as the nitrogen source, then washed with Gutnick medium and split into two cultures: one that was resuspended in Gutnick medium containing 15 mM ammonium and one in Gutnick medium without ammonium. Samples were then collected at 0, 15, 30, 60, and 150 min post-washout. For (**B**) and (**C**), total RNA (10 µg) was separated on an acrylamide gel and the transferred to a membrane, which was sequentially probed with labeled oligonucleotides for GlnZ and the 5S control RNA. (**D**) Growth of WT MG1655 and Δ*glnZ* (GSO1153) strains after nitrogen limitation. Cells were cultured in Gutnick medium with 0.4% glucose and 3 mM ammonium to stationary phase (OD_600_∼1.5). Cultures were then incubated another 48 h before subculturing to an OD_600_ of 0.05 into Gutnick medium with 0.4% glucose and 3 mM ammonium. After subculturing, OD_600_ was measured for 16 h. The average of four independent replicates is plotted. Error bars represent one standard deviation.

To see if we could detect the predicted products and determine when GlnZ levels are highest, we isolated total RNA from cells grown in several different carbon and nitrogen sources and examined GlnZ expression by northern analysis. For cells grown in defined Gutnick medium with 0.4% glucose or glycerol as the carbon source and 1.5 or 15 mM ammonium as the nitrogen source, GlnZ levels are highest when cells enter stationary phase (Figure 1B and Supplementary Figure S1). The most prominent band was ∼194 nt band, as seen previously (Kawano et al 2005) with minor bands of ∼174 and ∼150 nt (Figure 1B). The predicted ∼213 nt band was only detected with a long exposure (Supplementary Figure S1A). When cells were grown in LB or Gutnick medium with 0.4% *α*-ketoglutarate and 15 mM ammonium or 0.4% glucose and 15 mM glutamine, all GlnZ bands are significantly lower (Figure 1B and Supplementary Figure S1B).

With 15 mM glutamate as the nitrogen source, expression was highest in early log phase instead of in stationary phase (Figure 1B).

We also tested the effect of decreasing nitrogen by splitting a culture growing exponentially in Gutnick medium with 15 mM ammonium, washing the cells and then resuspending the cultures in Gutnick medium with 15 mM ammonium or in Gutnick medium lacking a nitrogen source. GlnZ levels increased rapidly for the cells transferred to the medium without nitrogen in contrast to the cells with sufficient nitrogen (Figure 1C). Together these observations indicate that GlnZ is induced by nitrogen starvation but is down regulated by *α*- ketoglutarate and glutamine.

### Δ*glnZ* mutants show decreased recovery from nitrogen starvation

Given the increased GlnZ levels under nitrogen limiting conditions, we compared the growth of WT and Δ*glnZ* cells exposed to long nitrogen starvation as described previously (34). Since *glnA* and *glnZ* share the same terminator, the *glnZ* deletion was constructed by replacing the 3’ UTR with sequence lacking homology to *E. coli* together with a modified *glnA* terminator maintaining the hairpin structure with a different nucleotide sequence. Immunoblot analysis documented that the *glnZ* deletion does not affect GlnA protein levels (Supplementary Figure S1C). The WT and Δ*glnZ* cultures were grown in Gutnick medium containing 0.4% glucose and 3 mM ammonium to stationary phase, where they reached nitrogen starvation. The cells were incubated in the nitrogen-starvation conditions for an additional 48 h, and then subcultured into fresh Gutnick medium containing 0.4% glucose and 3 mM ammonium. Time points taken over a 16 h period after subculturing showed a growth defect in the Δ*glnZ* strain compared to WT (Figure 1D).

These data indicate that GlnZ is required for optimal recovery from low nitrogen conditions.

### NtrC-mediated transcriptional regulation of GlnZ

Given the predicted GlnZ promoter (32), as well as the possibility of processing from the *glnA* transcript, we wanted to understand how GlnZ levels are influenced by transcriptional regulation. Due to the high levels of GlnZ seen in stationary phase, we first examined GlnZ levels in a strain lacking σ^S^, the stationary phase sigma factor encoded by *rpoS* (Supplementary Figure S2A). While GlnZ levels overall were lower in the Δ*rpoS* strain, we still observed induction pointing to only a minor contribution of σ^S^, especially compared to the known σ^S^-dependent sRNA SdsR (35). Previous studies (36, 37) have shown there are two promoters upstream of *glnA*, one of which is dependent on both NtrC and σ^54^ (Figure 1A). Thus, we next examined GlnZ levels in a strain lacking NtrC (Figure 2A) and found that GlnZ levels were strongly reduced in the Δ*ntrC* mutant. This regulation likely comes from NtrC activation of the *glnA* promoter as we did not detect GlnZ in a strain with a deletion of this upstream promoter (Figure 2A). Transcriptional regulation by NtrC also is consistent with the decreased levels of GlnZ seen for cells grown with *α*-ketoglutarate as the sole carbon source, as *α*-ketoglutarate is known to inhibit NtrC (38, 39).

**Figure 2.**
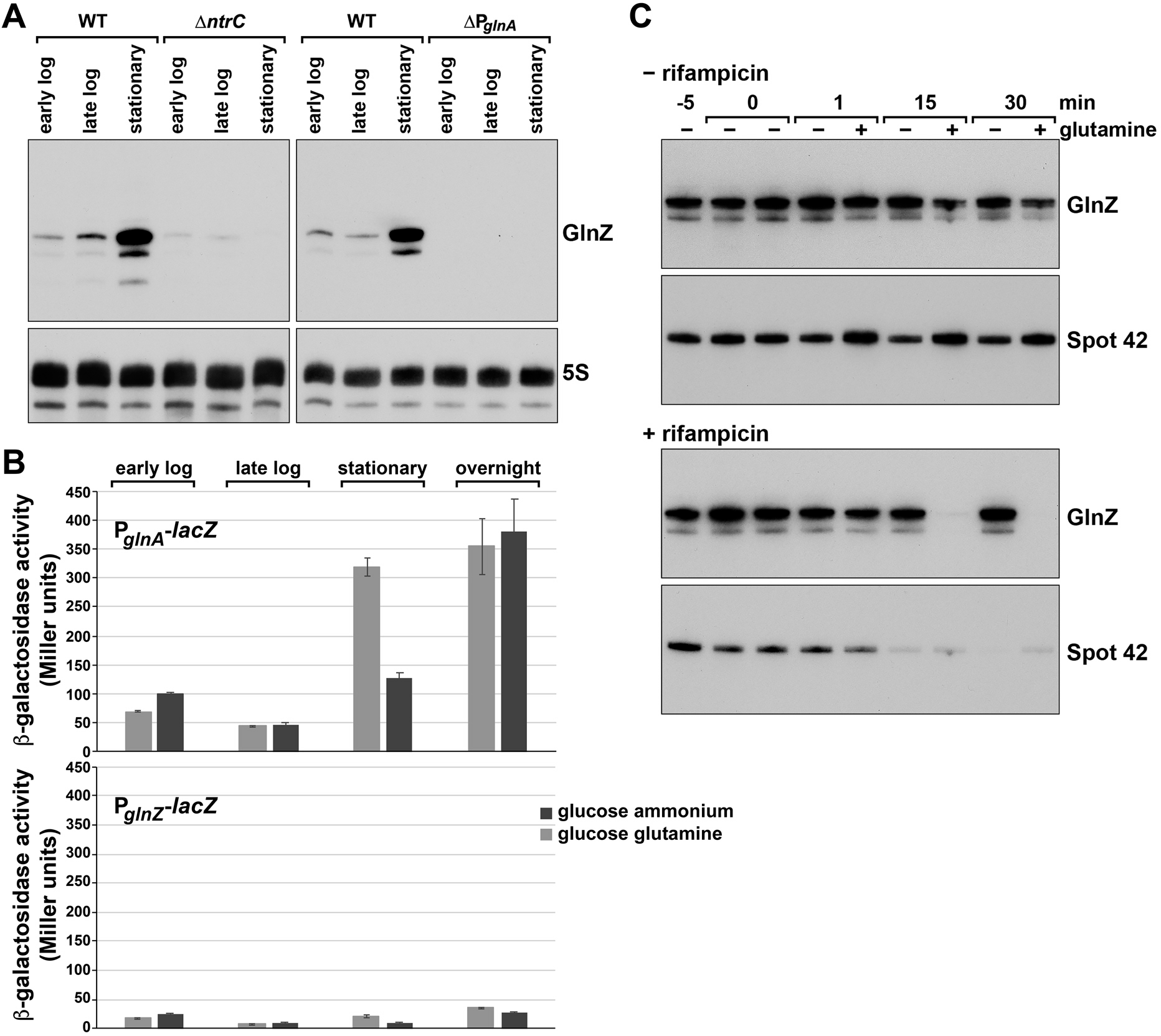
Transcriptional and post-transcriptional regulation of GlnZ levels. (**A**) Northern blot analysis of GlnZ levels in wt, Δ*ntrC* (GSO1151) and ΔP*_glnA_* (GSO1152) strains of *E. coli* cultured to stationary phase in Gutnick medium with 0.4% glucose and 15 mM ammonium and 1.5 mM glutamine. Northern analysis was carried out as in Figure 1. (**B**) *β*-galactosidase activity for PM1205 *P_glnA_-lacZ* (GSO1156) or *P_glnZ_-lacZ* (GSO1157) strains grown in Gutnick medium with 0.4% glucose and 15 mM ammonium or 15 mM glutamine. Samples were taken at early logarithmic, late logarithmic, stationary, and overnight phases of growth. The average of three independent replicates is plotted. Error bars represent one standard deviation. (**C**) Northern blot analysis of GlnZ levels in WT *crl_-_* MG1655 cells with and without treatment with rifampicin. A culture was grown to stationary phase (OD_600_∼3.5) and then split with one culture treated with rifampicin. After 5 min, cultures were split again, and 15 mM glutamine was added to one plus- rifampicin and one minus-rifampicin culture. Samples were taken at the indicated times and analyzed by northern analysis as in Figure 1.

To further investigate whether GlnZ is transcribed from its own promoter, we also generated chromosomal *lacZ* reporter fusions to P*_glnA_* and P*_glnZ_* at a heterologous site on the chromosome. *β*-galactosidase activity was measured for both strains in Gutnick medium with 0.4% glucose and 15 mM ammonium or 15 mM glutamine across growth. Consistent with the known induction of GlnA in response to nitrogen starvation, expression from P*_glnA_* was strongly elevated in stationary phase and overnight cultures (Figure 2B), particularly for cells grown in glutamine known to lead to starvation earlier in growth (40). In contrast, minimal *β*- galactosidase activity was detected for P*_glnZ_*-*lacZ* under all conditions tested. Together with the decreased GlnZ levels in the *glnA* promoter deletion strain, these data suggest that under these growth conditions, GlnZ is primarily processed from the *glnA* transcript, rather than transcribed from a promoter internal to *glnA*.

### Post-transcriptional regulation of GlnZ in response to glutamine

Given that growth in glutamine has been shown to activate the NtrC response (40), as demonstrated by activation of P*_glnA_*-*lacZ* in Figure 2B, we were surprised by the reduction in GlnZ levels for cells grown in glutamine. To test whether this could be due to post-transcriptional regulation, cells were grown to exponential phase in Gutnick medium with 15 mM ammonium and 0.4% glucose, then treated with the transcription inhibitor rifampicin.

Glutamine (15 mM) was then added to half of each culture. We detected lower levels of GlnZ in all samples exposed to glutamine, but this decrease was significantly greater when the cells were also treated with rifampicin (Figure 2C). The levels of a control sRNA, Spot 42, did not decrease with glutamine treatment.

To directly test whether the impact of glutamine is independent of the *glnA* and *glnZ* promoters, we cloned the 213 and 194 nt versions of GlnZ behind the P_lac_ promoter on the pNM46 vector, which also carries *lacI* (pBR-*lacI*). Expression from this plasmid is tightly regulated, with no RNA detected in the absence of IPTG inducer in a Δ*glnZ* background (Supplementary Figure S2B). We found that the levels of GlnZ transcribed from pBR-*lacI* again were lower in the presence of glutamine in cells treated with rifampicin (Supplementary Figure S2C). These observations show that glutamine reduces GlnZ levels by a post-transcriptional mechanism.

### GlnZ is significantly double stranded and cleaved by RNase III

To learn more about the GlnZ structure and possible post-transcriptional processing we probed in vitro synthesized GlnZ_194_ RNA with RNase III, which cleaves double-stranded RNA, and RNase T1, which cleaves 3’ of single-stranded G residues (Figure 3). This analysis showed extensive cleavage by RNase III, suggesting that long stretches of GlnZ are double-stranded, and allowed us to predict a structure for GlnZ. We suggest that the 5’ end can fold into two different alternative secondary structures (one shown in Figure 3A and the second forming between the underlined sequences). The first stem-loop depicted here could be an ideal substrate for cleavage by ribonuclease E, which cleaves single-stranded U-rich RNA followed by a 3’ stem-loop (41, 42), to give the∼174 nt form of GlnZ. The second stem-loop shown is consistent with one of the structures predicted for the repetitive extragenic palindromic (REP) sequence (43) found internal to GlnZ. While the functions of REP sequences in the genome are not well understood, they are known to give rise to highly-structured RNAs, consistent with the extensive RNase III cleavage in this portion of GlnZ.

**Figure 3.**
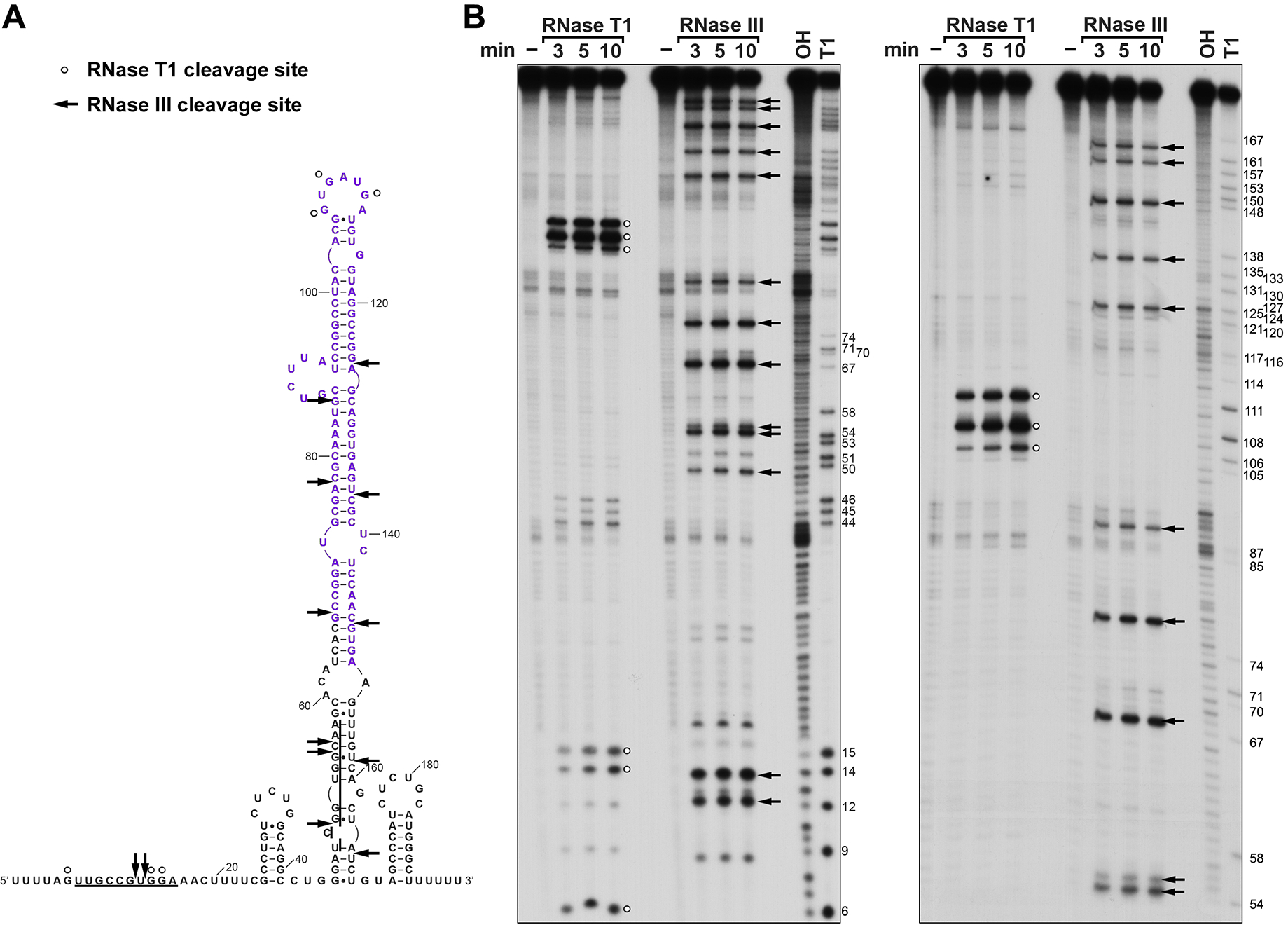
Predicted GlnZ structure. (**A**) Predicted secondary structure of GlnZ_194_ based on in vitro structure probing. Underlined sequences indicate location of possible alternative base pairing. Nucleotides corresponding to the REP sequence (64) are in purple font. (**B**) In vitro transcribed GlnZ_194_ was incubated with either water, RNase T1, or RNase III for 3, 5, or 10 min. Samples were then resolved, alongside an OH and T1 ladder, on a urea polyacrylamide gel for two lengths of time (left: 75 min, right: 150 min). For (**A**) and (**B**), white dots indicate cleavage by RNase T1, and arrows denote cleavage by RNase III.

Given the observed post-transcriptional regulation of GlnZ levels and cleavage of the long GlnZ stem by the double-stranded endonuclease RNase III in vitro, we wondered if RNase III impacted the downregulation of GlnZ in cells grown with glutamine. To test this, we assayed GlnZ levels in wild type and *rnc* mutant cells lacking RNase III, both strains treated with rifampicin and then incubated without and with glutamine (Supplementary Figure S3A).

Interestingly, GlnZ levels were decreased with and without glutamine in the *rnc* mutant cells. These observations suggest RNase III blocks an unknown factor that decreases GlnZ levels, and this positive effect of RNase III is not observed in the presence of glutamine. To determine whether the regulation by glutamine and RNase III are specific to GlnZ, we also probed the northern blots for Spot 42, not known to be digested by RNase III, and SdsR, reported to be cleaved by RNase III (44). Glutamine did not lower the levels of the Spot 42 RNA (Figure 2C and Supplementary Figures S2C and S3A). However, as for GlnZ, we observed significantly lower SdsR levels in the presence of glutamine or the absence of RNase III (Supplementary Figures S2C and S3A). Thus, we suggest a common, unidentified factor modulates GlnZ and SdsR levels. We did not observe significant effects of either 15 mM or 50 mM glutamine on RNase III-dependent cleavage of GlnZ_194_ RNA in vitro (Supplementary Figure S3B), consistent with the conclusion that another factor is required.

### Putative GlnZ targets identified by RIL-seq and MAPS approaches

To understand the mechanism by which GlnZ contributes to recovery from nitrogen starvation, we next examined the regulatory role of GlnZ. The previous observations that GlnZ levels decrease in a *Δhfq* mutant strain (16) and GlnZ co-purifies with Hfq (45, 46), suggested that GlnZ acts as an Hfq-dependent base pairing RNA. Consistent with this suggestion, a number of putative targets were detected as chimeras with GlnZ by RNA Interaction by Ligation and sequencing (RIL-seq) for cells grown in LB, M63, and iron limited media (45, 46) (Supplementary Table S4).

Given the importance of GlnZ after nitrogen starvation, we also wanted to look for putative GlnZ targets in cells grown to stationary phase in Gutnick medium with ammonium. Thus, we tagged the 5’ end of GlnZ with the MS2 sequence to carry out MS2-affinity purification coupled with RNA sequencing (MAPS) for cells grown with limited nitrogen.

Northern analysis confirmed that MS2-GlnZ_194_ was produced stably and was enriched in the eluate compared to the lysate (Supplementary Figure S4A). As expected, Hfq co-purified with MS2-GlnZ_194_ (Supplementary Figure S4A). Sequencing of the co-purifying RNAs revealed enrichment of numerous putative targets with MS2-GlnZ_194_ (Supplementary Table S5).

Interestingly, many highly-enriched mRNAs also were represented in abundant chimeras in the RIL-seq experiments performed under different conditions, though some unique candidate targets were identified by each protocol (Supplementary Tables S4 and S5). These unique candidates may be due in part to differences in gene expression given the different growth conditions. Browser images of the sequencing data for the target candidates enriched most strongly in both datasets are shown in Figure 4 and Supplementary Figure S4, together with examples that were enriched in only the RIL-seq (*rdcA* (*yjdA*)-*rdcB* (*yjcZ*)) or MAPS (*yahO*) data, respectively.

**Figure 4.**
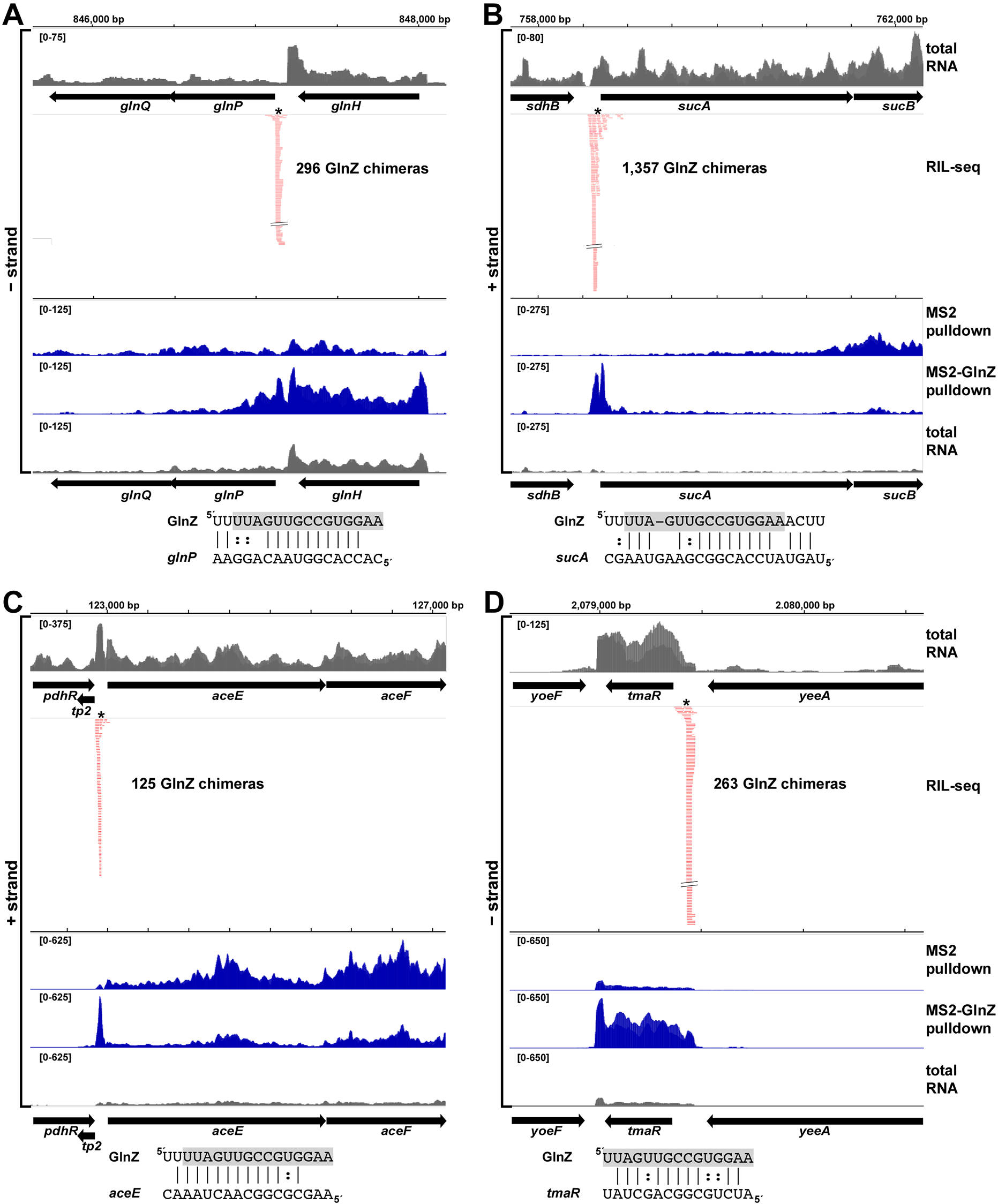
GlnZ interactions with (**A**) *glnP*, (**B**) *sucA*, (**C**) *aceE*, and (**D**) *tmaR* as detected by RIL-seq (46) or MAPS. In browser images, total RNA is in grey, RIL-seq chimeras detected in M63 medium are in red, and MAPS data for cultures grown in Gutnick medium with 0.04% glucose and 15 mM ammonium at stationary phase is in blue for the pulldown with either MS2 or MS2-GlnZ overexpression. Predicted regions of interaction with GlnZ are depicted below each browser image. The GlnZ seed region is highlighted in grey. The position of the predicted interaction is indicated by an asterisk in the browser images.

For *sucA*, encoding a component of the *α*-ketoglutarate dehydrogenase complex, and *aceE*, encoding pyruvate dehydrogenase, only the 5’ UTR region was enriched in both approaches. For other examples such as *glnH-glnP*, encoding components of a glutamine ABC transporter, *tmaR* (*yeeX*), encoding a regulator of the PTS system, *fre*, encoding NAD(P)H-flavin reductase and McaS, a small base pairing RNA, transcripts corresponding to the full gene were enriched by MAPS though the interaction detected by RIL-seq was very defined. The reasons for the more extensive MAPS signals are not known but may be due to different GlnZ mechanisms of action or differences in the processing of the target RNAs.

Previous RIL-seq analysis predicted that the region between 194 nt and 174 nt from the GlnZ 3’ end encompasses a seed sequence for GlnZ base pairing with targets (45). Possible base pairing between this seed region and the targets could be predicted in the sequences overlapping the RIL-seq signals (Figure 4 and Supplementary Figure S4). We noted that the position of the RIL-seq signal and predicted base pairing varied significantly between the targets. Some base pairing regions overlapped or were adjacent to the RBS (*sucA*, *glnP* and *rdcB*), as is commonly observed for sRNA regulation, while others were found much farther upstream of the RBS (60- 80 nt) (*aceE* and *tmaR*) or in the 3’ UTR (*fre*), suggesting more complex mechanisms of regulation. We first examined the effects of GlnZ on *sucA* and *glnP*, given their roles in carbon and nitrogen metabolism and the predicted base-pairing near the RBS.

### GlnZ represses *glnP* and *sucA* expression to affect nitrogen flux

Direct GlnZ pairing to *glnP* (Figure 5A) and *sucA* (Figure 5B) was tested by examining the effects of WT and mutant GlnZ_194_ overexpression on fusions to these genes. Compared to the empty vector control, overexpression of GlnZ_194_ led to a decrease in mCherry fluorescence (Figure 5C) for a chromosomal translational fusion between the *glnP* 5’ UTR encompassing the predicted target site and mCherry. GlnZ_194_-M1 overexpression did not repress the *glnP*-mCherry reporter. In contrast, a *glnP*-mCherry reporter carrying a mutation compensatory to GlnZ_194_-M1 was repressed by GlnZ_194_-M1 rather than GlnZ_194_ although the mutant fusion had lower levels of expression. Overall, these data demonstrate that GlnZ represses *glnP* through direct base-pairing.

**Figure 5.**
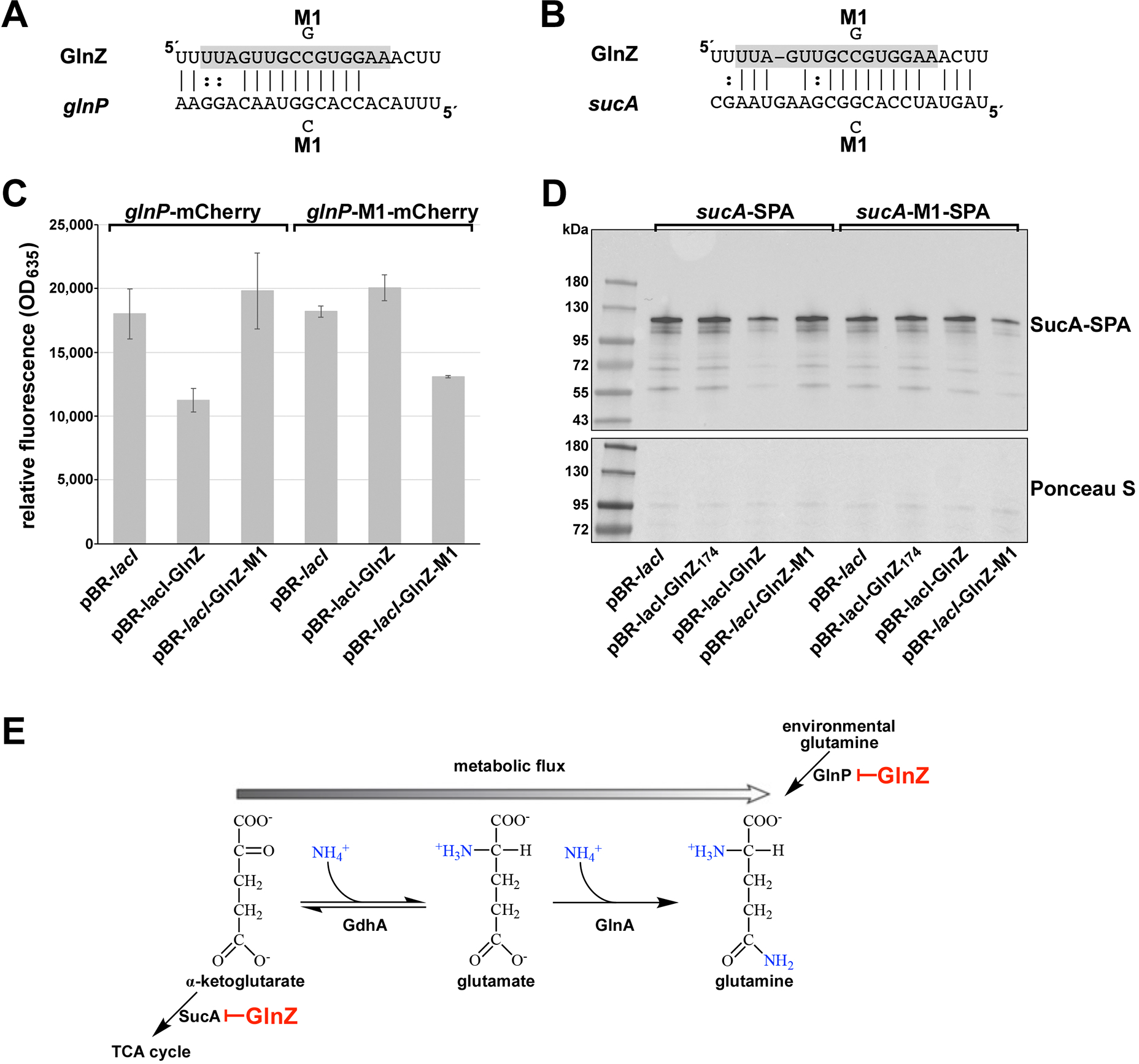
GlnZ downregulation of *sucA* and *glnP*. (**A**) Predicted base pairing between GlnZ and *glnP* 5’ UTR and mutations generated. (**B**) Predicted base pairing between GlnZ and *sucA* 5’ UTR and mutations generated. For (**A**) and (**B**), GlnZ seed region is highlighted in grey. (**C**) GlnZ repression of a *glnP-*mCherry fusion. The indicated strains were grown for 3 h in Gutnick medium with 0.4% glucose and 15 mM ammonium and 1 mM IPTG to induce WT and mutant GlnZ expression and either 0.2% arabinose (*glnP*-mCherry) or 2% arabinose (*glnP-*M1- mCherry) to induce the mCherry fusion. Relative fluorescence units were determined by measuring OD_635_ for each sample and normalizing by OD_600_. The average of three independent replicates are plotted. Error bars indicate one standard deviation. (**D**) Immunoblot analysis of SucA-SPA levels. The indicated strains were grown to mid logarithmic phase (OD_600_ ∼1.0) in Gutnick medium with 0.4% glucose and 15 mM ammonium and 1 mM IPTG to induce WT and mutant GlnZ expression. Samples were normalized by OD_600_, separated by SDS-PAGE, and analyzed using immunoblot analysis. The Ponceau S-stained membrane serves as the loading control. (**E**) Model of GlnZ effects on carbon and nitrogen metabolism. Carbon and nitrogen metabolism in *E. coli* are interconnected by the interconversion of *α*-ketoglutarate, glutamate and glutamine through the addition or removal of ammonium. Amino groups added to glutamate or glutamine after each step are highlighted in blue. Arrows indicate the flow of substrates to products with a double headed arrow indicating conversion in both directions. Enzymes involved in the reaction are listed below the arrows.

As a *sucA-*mCherry reporter fusion produced only low levels of mCherry expression, we examined the effects of GlnZ overexpression on SucA-SPA protein levels. Western analysis showed that upon overexpression of GlnZ_194_, SucA-SPA levels decrease compared to a pBR-*lacI* vector control. As expected from the position of the seed sequence between nucleotides 194 and 174, SucA-SPA levels are unaffected by the overexpression of GlnZ_174_ (Figure 5D). The GlnZ_194_-M1 seed sequence mutant also did not repress SucA-SPA levels, though repression was restored for a *sucA-M1-SPA* fusion indicating this regulation occurs through direct base pairing. We did note that *sucA-M1-SPA* also was repressed by WT GlnZ_194_ overexpression in comparison to a pBR-*lacI* vector control, indicating that the single point mutation does not eliminate regulation in the context of the *sucA*-*SPA* fusion.

The GlnP and SucA proteins both play roles in glutamine accumulation (Figure 5E).

Consistent with the strong regulation of *α*-ketoglutarate dehydrogenase levels, cells with GlnZ overexpression from pBR-GlnZ_194_ showed reduced growth in medium with *α*-ketoglutarate as the sole carbon source with either ammonium or glutamine as the nitrogen source (Supplementary Figure S6A). We propose that under conditions of limiting nitrogen, particularly limiting glutamine, regulation by GlnZ promotes accumulation of this nitrogen sink.

### Varied regulatory mechanisms affect other GlnZ targets with varied physiological roles

While GlnZ base pairing occurs close to the RBS in the 5’ UTR of *sucA* and *glnP*, the positions of the pairing sites predicted for several other GlnZ targets are less canonical. For both *aceE* and *tmaR*, the predicted base pairing is significantly upstream of the RBS (Supplementary Figure S5). In the case of *aceE*, several shorter transcripts can be detected for the 5’ UTR (Supplementary Figure S6B), possibly reflective of extensive regulation in the leader region including RNase III cleavage (47). The levels of all the shorter transcripts decrease with GlnZ overexpression (Supplemental Figure S6B) suggesting GlnZ might facilitate or interfere with the upstream regulation. Little is known about the *tmaR* transcript, and we did not observe any changes in the levels of a TmaR-YFP fusion upon GlnZ expression (Supplementary Figure S6C). GlnZ is predicted to base pair upstream of the reported 5’ end of the predominant McaS transcript. However, longer forms of McaS were previously suggested (48), and northern analysis demonstrated that larger McaS transcripts are present at low levels and the abundance of these longer transcripts is decreased by GlnZ overexpression (Supplementary Figure S6B). An additional band also appears upon GlnZ overexpression, potentially corresponding to a cleavage product formed due to regulation by GlnZ. The predicted pairing with the *fre* transcript is an extensive sequence in the 3’ UTR. We did not characterize this pairing further but do not think the *fre* 3’ UTR is serving as an sRNA since this region is the first RNA in all the RIL-seq chimeras, a feature of mRNA targets.

### Some GlnZ-mediated repression occurs through RNase III cleavage

Given that *sucA* and *aceE* are known targets of RNase III (47, 49), we wondered whether GlnZ down regulation of these targets could be due to cleavage by RNase III. Thus, we examined the effect of GlnZ overexpression on SucA-SPA protein levels in a Δ*rnc* background (Figure 6A). In this mutant strain, SucA-SPA levels remain constant upon GlnZ_213_ or GlnZ_194_ overexpression indicating GlnZ repression of SucA-SPA is dependent on RNase III. In contrast, downregulation by RyhB, another sRNA known to repress the *sdh-suc* operon (Masse and Gottesman 2002), was unaltered in the Δ*rnc* mutant. By primer extension analysis, we also detected a position of increased GlnZ_194_-dependent RNase III cleavage of *aceE* 65 nt downstream of the predicted base pairing (Figure 6B). Whether the base pairing leads to a change in *aceE* 5’ UTR structure to promote cleavage is not known, but our observations are consistent with GlnZ directing RNase III cleavage at some targets.

**Figure 6.**
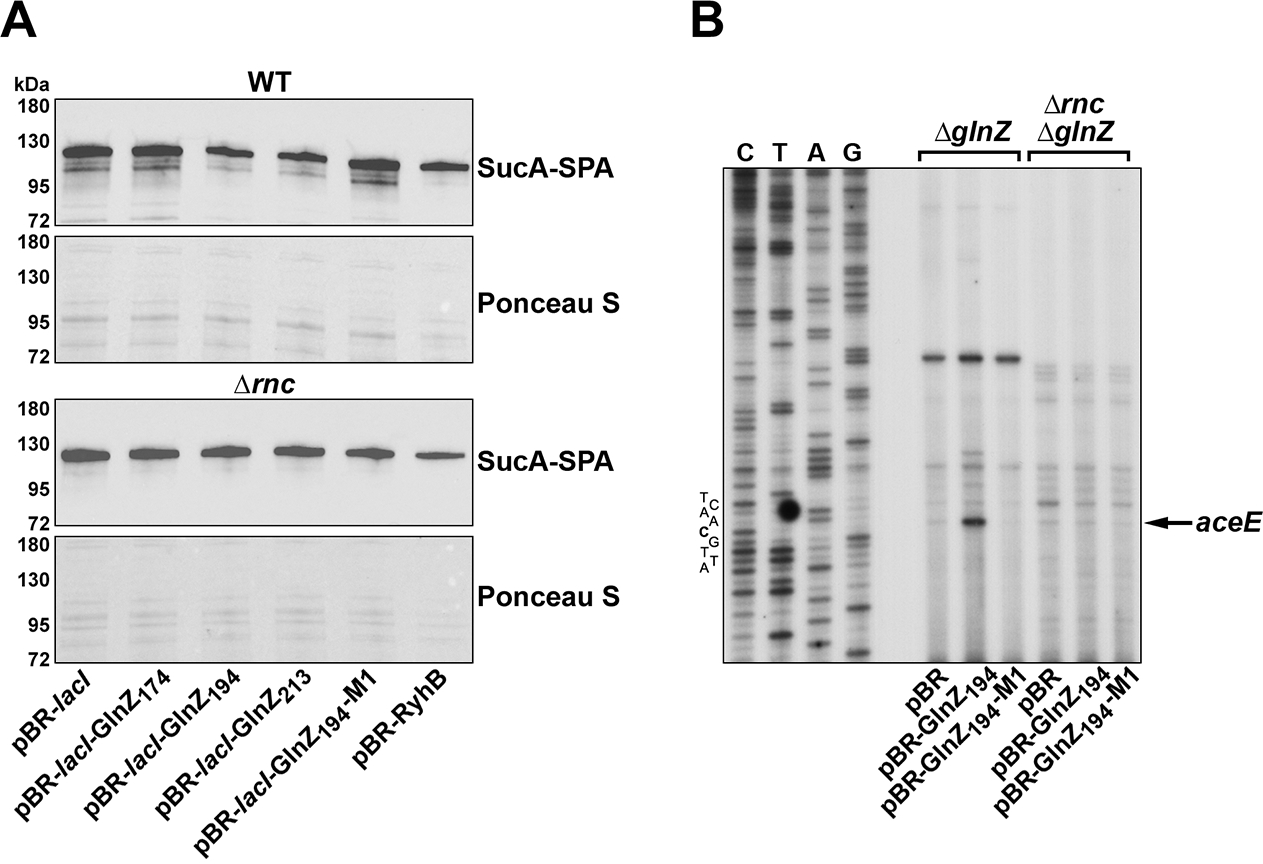
Direct GlnZ base pairing leads to RNase III-dependent cleavage. (**A**) WT MG1655 or Δ*rnc* (GSO1154) strains carrying *sucA-SPA* and harboring the indicated plasmids were grown in Gutnick medium with 0.4% glucose and 15 mM ammonium and 1 mM IPTG to induce WT or mutant GlnZ expression. Once cells reached mid-logarithmic phase (OD_600_ ∼1.0), samples were normalized by OD_600_, separated by SDS-PAGE and analyzed using immunoblot analysis. The Ponceau S-stained membranes serve as loading controls. (**B**) Primer extension analysis of *aceE* was performed on RNA isolated from Δ*glnZ* (GSO1153) and Δ*glnZ* Δ*rn*c (GSO1155) strains harboring indicated plasmids and grown to late logarithmic phase (OD_600_ ∼ 1.5) in Gutnick medium with 0.4% glucose and 15 mM ammonium. WT or mutant GlnZ expression was induced with 1 mM IPTG. A sequencing ladder is shown for *aceE*.

### GlnZ sequence varies across bacterial species

Although the full GlnZ sequence is not broadly conserved, sRNAs can be detected downstream of the glutamine synthetase gene in RNA-seq data for *Salmonella enterica* (50) and *Klebsiella* (M. Buck, personal communication). Additionally, during the course of this work, an sRNA denoted EsrF was detected downstream of *glnA* in the EHEC strain O157:H7 (51). To explore the evolutionary relationship between these RNAs, we investigated the 3’ UTR of the conserved glutamine synthetase gene in other bacterial species. This analysis revealed that the overall sequences had different lengths and were not conserved (Supplementary Figures S7A, S8A).

However, all of the 3’ UTRs contain the ∼20 nt seed sequence region of GlnZ, which typically is the most highly conserved region of bacterial sRNAs (52) as well as a conserved terminator (Supplementary Figures S7, S8A). These putative sRNAs can be categorized into three classes: (I) sRNAs whose seed sequence is located directly upstream of the terminator, with very few nucleotides between the seed region and the *glnA* stop codon (EHEC O157:H7 EsrF), (II) sRNAs with a seed sequence directly upstream of the 3’ terminator but with a significant number of nucleotides between the seed region and the *glnA* stop codon (*S. enterica* GlnZ), and (III) sRNAs where the seed sequence is located at the 5’ end near the *glnA* stop codon with a large REP sequence insertion between the seed sequence and the conserved terminator stem-loop (*E. coli* GlnZ) (Figure 7A, Supplementary Figure S7A). These classes of GlnZ sRNAs group by phylogeny (Figure 7B). We suggest that bacteria initially contained the minimal Class I GlnZ sequence. However, over time, large segments of DNA appear to have been inserted either upstream or downstream of the conserved seed sequence. Despite these large insertions, all GlnZ homologs have similar linear distances between the seed sequence and terminator regions, given that the REP sequence inserted in the *E. coli* Class III RNA forms a long stem-loop structure (Figure 3).

**Figure 7.**
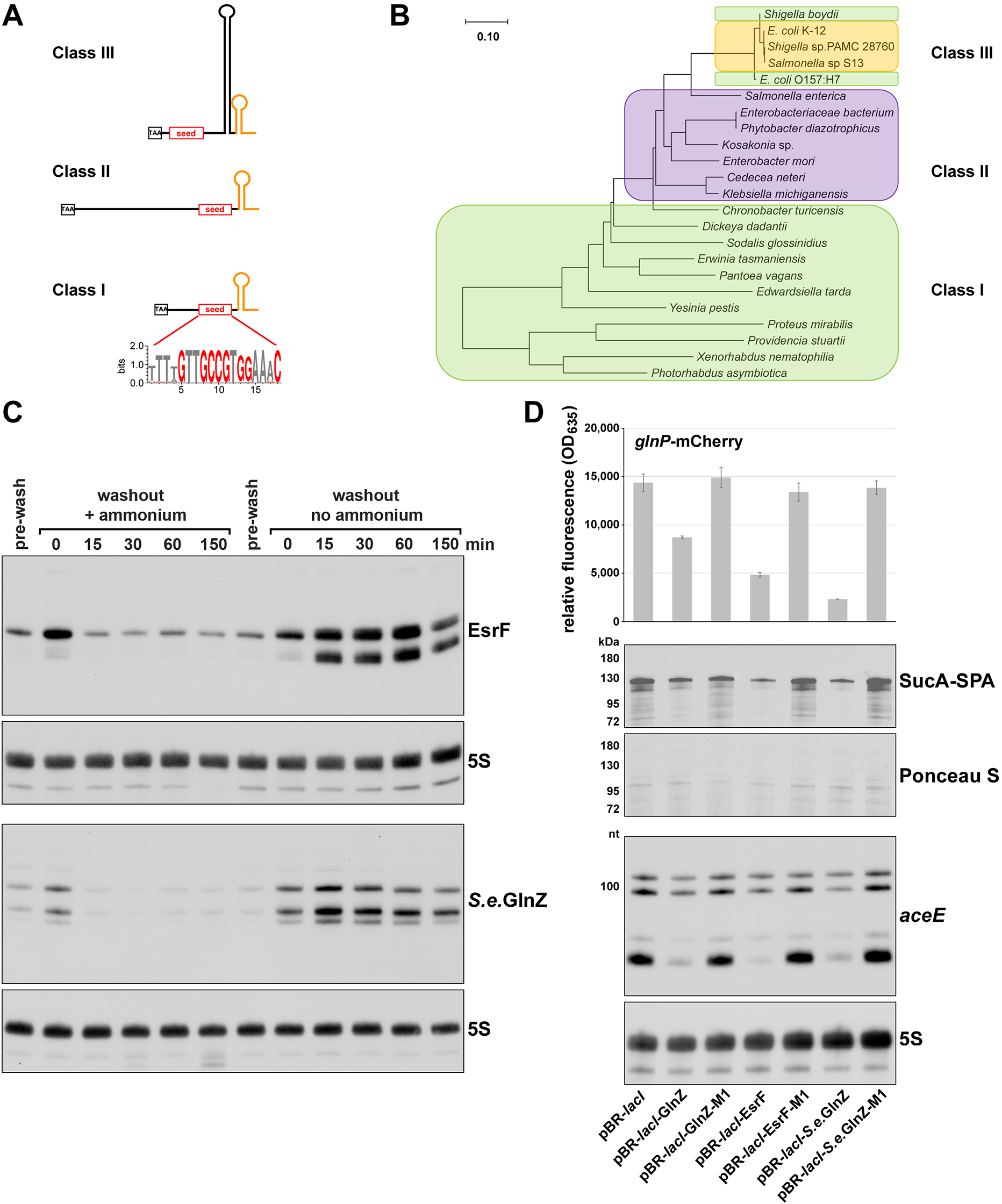
sRNAs encoded downstream of *glnA* in EHEC and *S. enterica* have insertions but are induced by low nitrogen and repress similar targets. (**A**) Diagram depicting the three classes of GlnZ transcripts. The stop codon of *glnA* is indicated with the black box, the conserved GlnZ seed region is in red font and the conserved terminator is in orange. Logo depicts the conservation of the GlnZ seed region in the Gammaproteobacteria indicated in Supplementary Figure S7B. (**B**) Phylogenetic tree of Gammaproteobacteria containing a GlnZ homolog. The evolutionary history was inferred as described in Materials and Methods, and the tree with the highest log likelihood (-27049.09) is shown. The tree is drawn to scale, with branch lengths measured in the number of substitutions per site. The three classes of GlnZ transcripts are indicated by colored boxes: Class I in green, Class II in purple, and Class III in yellow. (**C**) EHEC O157:H7 EsrF and *S. enterica* GlnZ are induced by low nitrogen. EHEC O157:H7 and *S. enterica* cells were grown and analyzed by northern blots as described in Figure 1C. The top membrane was probed for EsrF and 5S, and the bottom membrane was probed for *S.e.* GlnZ and 5S. (**D**) EHEC O157:H7 EsrF and *S. enterica* GlnZ repress GlnZ targets *glnP*, *sucA* and *aceE* in *E. coli* K-12. Relative fluorescence levels for the chromosomal *glnP-mCherry* fusion strain transformed with plasmids overexpressing *E. coli* K-12 GlnZ, EHEC EsrF or *S. enterica* LT2 GlnZ and the corresponding M1 mutants. mCherry fluorescence was measures as in Figure 5C. The average of three replicates is shown, and error bars represent one standard deviation. Immunoblot analysis of SucA-SPA levels was carried out as in Figure 5D. Northern analysis was performed using Δ*glnZ* (GSO1153) strains harboring the indicated plasmids were grown in Gutnick medium with 0.4% glucose and 15 mM ammonium and 1 mM IPTG. The blot was probed with labeled oligonucleotides for the *aceE* 5’ UTR and 5S RNA.

### Low nitrogen also induces expression of GlnZ homologs

In their characterization of EsrF, Jia et al. reported a slight increase in EsrF level in high ammonium compared to low ammonium for cells grown in M9 minimal medium (51). These observations led Jia et al. to conclude that EsrF is induced by high levels of ammonium, which is opposite to our conclusion that GlnZ is induced by low nitrogen conditions. To directly examine the levels of the GlnZ homologs in low and high nitrogen, the EHEC O157:H7 EDL933 and *S. enterica* LT2 strains were grown in Gutnick medium containing 0.4% glucose and 15 mM ammonium and samples were collected across multiple phases of growth. Northern analysis showed that EsrF and *S. enterica* GlnZ (*S.e.* GlnZ) levels are highest in stationary phase, like *E. coli* K-12 GlnZ levels (Supplementary Figure S8B and S8C). Additionally, when EHEC O157:H7 or *S. enterica* cultures were deprived of nitrogen, EsrF and *S.e.*GlnZ levels also were rapidly induced (Figure 7C). Thus, EsrF and *S.e.*GlnZ are expressed under the same conditions as *E. coli* GlnZ.

### GlnZ with varied sequences repress *E. coli* targets

To determine if EHEC EsrF and *S.e.*GlnZ could regulate *E. coli glnP*, *sucA* and *aceE* similar to *E. coli* GlnZ, we compared the effects of ectopically overexpressing these sRNAs in *E. coli* along with the corresponding seed region mutants (M1) on target fusions or transcripts (Figure 7D). Upon overexpression of EsrF or *S.e.*GlnZ, GlnP-mCherry levels decrease dramatically compared to the vector control and the EsrF-M1 or *S.e.*GlnZ-M1 seed sequence mutants (Figure 7D). EsrF-M1 or *S.e.*GlnZ-M1 were able to repress the GlnP-M1-mCherry compensatory mutant (Supplementary Figures S8D and S8E), indicating direct base pairing. The effects of EsrF and *S.e.*GlnZ overexpression on SucA-SPA protein and *aceE* 5’ UTR levels were similar; WT versions of the sRNAs decreased the levels while the mutant derivatives did not. These results demonstrate that despite a lack of overall sequence similarity, EsrF and *S.e.*GlnZ regulate *glnP*, *sucA* and *aceE* like *E. coli* GlnZ, consistent with the conserved seed sequence (Supplementary Figure S8F).

## DISCUSSION

A previous cloning-based screen to identify sRNAs in *E. coli* identified a sRNA at the 3’ end of the *glnA* open reading frame (16). Renamed GlnZ, the sRNA has multiple forms with different 5’ ends and a single 3’ end, though the predominant form is 194 nt. We showed that GlnZ expression is elevated under conditions of low nitrogen and that the sRNA enhances recovery from low nitrogen conditions. Multiple GlnZ targets were identified by RIL-seq and MAPS analysis, of which *sucA* and *glnP,* impact nitrogen flux, while other targets likely impact other aspects of metabolism. Finally, analysis of GlnZ conservation demonstrated that the seed sequence is highly conserved across Enteric bacterial species, despite a lack of overall sequence homology.

### GlnZ is induced by nitrogen limitation

GlnZ levels strongly increase upon nitrogen depletion (Figure 1C). Most of this induction is through transcriptional regulation of the *glnA* promoter by NtrC, as little GlnZ is detected in a strain lacking the response regulator (Figure 2A). Despite the predicted internal promoter at the 3’ end of *glnA*, no GlnZ was detected when the upstream *glnA* promoter was deleted, suggesting that under the conditions tested, GlnZ is primarily generated by processing from the NtrC- induced *glnA* transcript (Figure 2B). Whether there are other conditions that lead to induction of the internal promoter remains to be determined.

NtrC-mediated regulation of genes involved in nitrogen metabolism, biosynthesis or transport has been well-documented and typically occurs in response to either changes in nitrogen concentration or glutamine availability (3, 40). In contrast to what is observed for other NtrC-dependent genes, GlnZ levels are lower in the presence of glutamine. This glutamine-dependent regulation was found to be a post-transcriptional effect dependent on RNase III (Figure 2C). Interestingly, increased sRNA levels in the presence of RNase III and the absence of glutamine were also observed for other known sRNA targets of RNase III, demonstrating this regulation is not specific to GlnZ. RNase III activity is known to be positively regulated by phosphorylation by a bacteriophage T7 protein kinase (53) and negatively regulated by binding to the *E. coli* protein YmdB (54). The possibilities that glutamine causes changes in RNase III activity, or that the RNase III- and glutamine-dependent changes in sRNA levels occur via another factor such as a common base pairing target, warrant further investigation.

### GlnZ alters metabolism

Under conditions of limited nitrogen, *E. coli* alters its metabolism in response to the availability of ammonium or glutamine. Our data show that when ammonium is depleted, GlnZ levels increase, and the sRNA negatively regulates *sucA* and *glnP* (Figure 5C). Since GlnZ is downregulated by glutamine, it is likely when GlnZ levels are highest, the extracellular environment has no or low levels of glutamine. Under low glutamine conditions, GlnP would be upregulated transcriptionally by NtrC as part of the nitrogen starvation response (3). However, it would be energetically wasteful for the cell to continue to produce a glutamine transporter in the absence of extracellular glutamine; thus, downregulation by GlnZ would prevent GlnP translation. Downregulation of SucA should alter metabolic flux by channeling *α*-ketoglutarate from the TCA cycle into glutamate and glutamine, which are the major nitrogen sinks in the cell.

Other GlnZ targets identified by RIL-seq and MAPS play varied roles in metabolism. AceE is a subunit of the pyruvate dehydrogenase enzyme connecting glycolysis to the TCA cycle while Fre is a NAD(P)H-flavin reductase, which helps reactivate ribonucleotide reductase. While little is known about TmaR, this protein affects polar localization of enzymes of the phosphotransferase system (PTS) (55). Hfq can form polar foci under conditions of nitrogen starvation (56), raising the question of whether there is a connection between this phenomenon and the interaction between the GlnZ and *tmaR* RNAs. Conceivably further understanding of the GlnZ role in the regulation of these targets could give insights into the physiological response to low ammonium.

The mechanistic consequences of GlnZ base pairing with *aceE* 5’ UTR, the full-length *tmaR* transcript, longer versions of the McaS sRNA and the 3’ of *fre* along with the interaction between GlnZ and RNase III could also be subjects of future studies. Based on northern analysis (Supplementary Figure 6B), short transcripts corresponding to the 5’ UTR of *aceE* can be detected, suggesting additional regulation in this region. The levels of all these shorter transcripts are decreased by GlnZ overexpression consistent with the GlnZ- and RNase III-dependent cleavage we observe (Figure 6B).

### GlnZ-like sRNAs show limited overall conservation but similar regulation and targets

Maintaining nitrogen homeostasis is critical to all bacteria and many regulatory aspects are conserved. The NtrBC nitrogen stress response regulators are found across enteric bacteria. Additionally, other transcriptional regulators of nitrogen homeostasis have been identified in diverse bacterial species (57). Given the importance of nitrogen to cell viability, one can imagine that there is strong selective pressure to optimally regulate nitrogen flux at all levels including sRNA-mediated post-transcriptional regulation. Glutamine synthetase is integral to the nitrogen starvation responses and is found in all bacterial species (58). Thus, it is unsurprising that GlnZ- like sRNAs with similar regulatory functions are generated from multiple *glnA* mRNAs.

While evolution of sRNAs has not been studied extensively, it has been suggested that insertion of seed sequences, previously found to be the most conserved region (52, 59), into a 3’ UTR is a rapid way to evolve new sRNA regulators (reviewed in (60)). Consistent with this, the GlnZ seed sequence and terminator showed the greatest homology across bacterial species (Figure 7, Supplementary Figure S7A). Given the conservation of the seed sequence, we hypothesize that many mRNA targets are also conserved. Analysis of GlnZ targets in *E. coli* K- 12, EHEC, and *S. enterica* in this study, and by Miyakoshi et al., support this hypothesis documenting regulation of *sucA* and *glnP* by the homologs (61). Additionally, recent RIL-seq analysis of Hfq-binding sRNAs in enteropathogenic *E. coli* (EPEC) identified a GlnZ sRNA derived from the 3’ UTR of *glnA* and predicted *sucA* and *glnP* as potential binding partners of GlnZ in EPEC (62). Together, these data show that despite insertions in GlnZ across species, the conserved seed sequence allows for regulation of the same targets, providing a similar functional role.

Insertion of sequences up- or downstream of the GlnZ seed sequence might be expected to have significant effects on the sRNA structure and its ability to regulate its targets. However, structure probing data demonstrates that the REP sequence inserted between the seed sequence and terminator of GlnZ from *E. coli* K-12 is highly double-stranded, forming a long stem-loop structure (Figure 3). This leaves the seed sequence accessible for base pairing, while also maintaining a similar linear distance between the seed region and the terminator loop to that in the other GlnZ classes lacking this insertion. Insertion of a REP sequence, not present in other MgrR homologs, was noted in the MgrR sRNA in *Escherichia fergusonii* (63). In this case, a known MgrR target in *E. coli* was not regulated by *E. fergusonii* MgrR, despite identical seed sequences. It will be interesting to see how insertions and deletions affect the target specificity of other sRNAs as more and more homologs or paralogs are characterized across bacterial species.

In summary, the characterization of GlnZ provided insights into GlnZ expression, its physiological role in modulating carbon and nitrogen metabolic flux, and its functional conservation among Enteric bacteria. However, this work also revealed new questions that will form the basis for future studies of the response to limited nitrogen, mechanisms of sRNA- and RNase III-mediated regulation, as well as sRNA evolution.

## DATA AVAILABILITY

The NGS data discussed in this publication are accessible through GEO Series accession number GSE199845.

## ACKNOWLEDGEMENTS

We thank R. Levine for the kind gift of anti-glutamine synthetase antiserum and members of the Storz lab, M. Buck, S. Gottesman, M. Maurizi, S. Wigneshweraraj for helpful discussions.

## SUPPLEMENTARY DATA

Supplementary Data are available.

