## Supplementary for "A 3’ UTR-derived small RNA connecting nitrogen and carbon metabolism in enteric bacteria"

### Supplementary Data

### Supplementary Figures

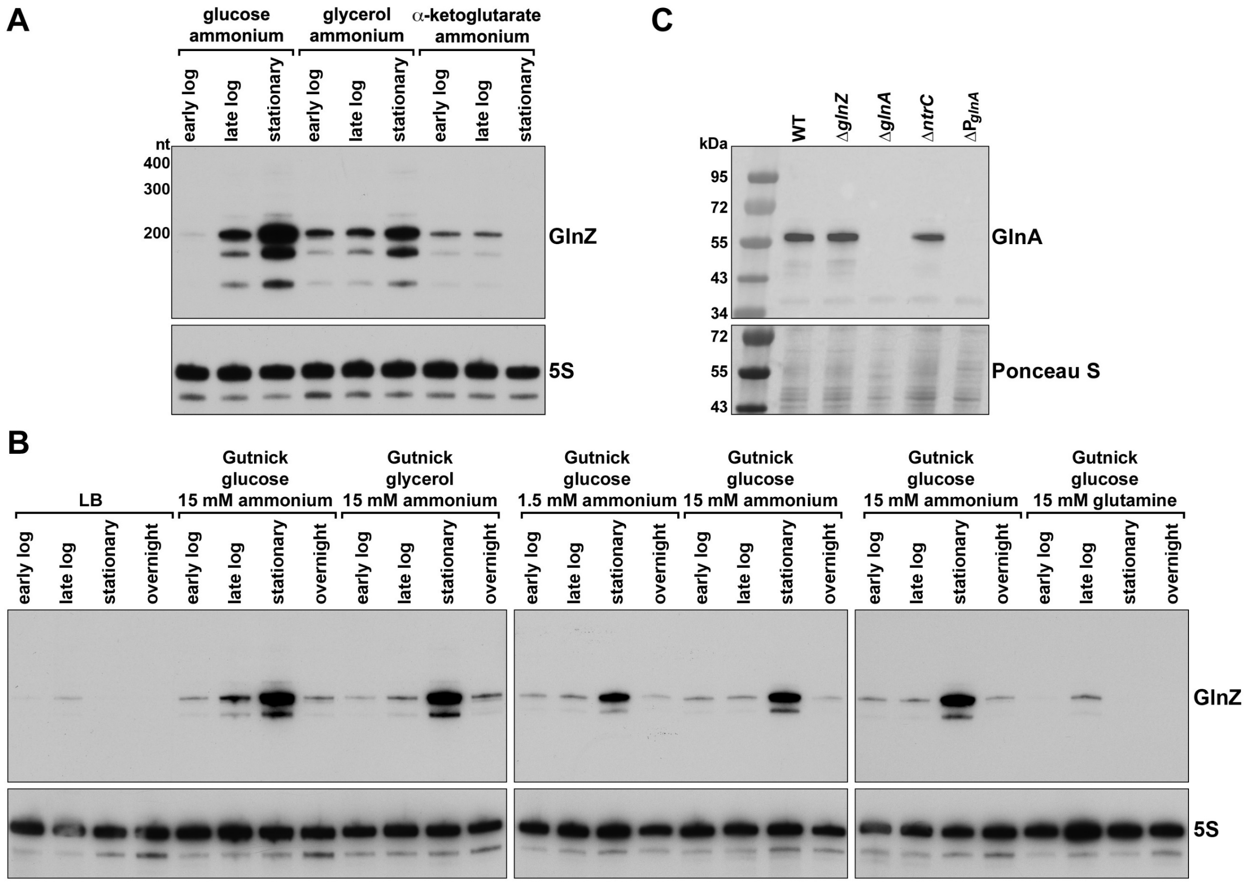

**Figure S1.** GlnZ expression across growth in different carbon and nitrogen sources. (**A**) Longer exposure of northern blot shown in Figure 1B. WT MG1655 was cultured to early logarithmic, late logarithmic or stationary phases in Gutnick medium with 0.4% glucose, glycerol or α-ketoglutarate as the sole carbon source and 15 mM ammonium as the nitrogen source. (**B**) Northern blot analysis of GlnZ levels with WT *crl*⁻ MG1655 (GSO983) cultured to early logarithmic ($OD_{600}$~0.2), late logarithmic ($OD_{600}$~0.7), stationary phase ($OD_{600}$~3.5) or overnight ($OD_{600}$~3.5) in LB or Gutnick medium with 0.4% glucose or glycerol as the sole carbon source and 1.5 or 15 mM ammonium or 15 mM glutamine as the nitrogen source. Total

RNA was collected and subjected to northern blot analysis with labeled oligonucleotide probes to GlnZ and 5S. (**C**) Immunoblot analysis of glutamine synthetase levels. The indicated strains were grown to stationary phase ($OD_{600}$ ~3.5) in Gutnick medium with 0.4% glucose and 15 mM ammonium and 1.5 mM glutamine (to allow growth of the Δ*glnA* and $\Delta P_{glnA}$ strains). Samples were normalized by $OD_{600}$, separated by SDS-PAGE, and analyzed using immunoblot analysis with an anti-glutamine synthetase antibody. The Ponceau S-stained membrane serves as the loading control.

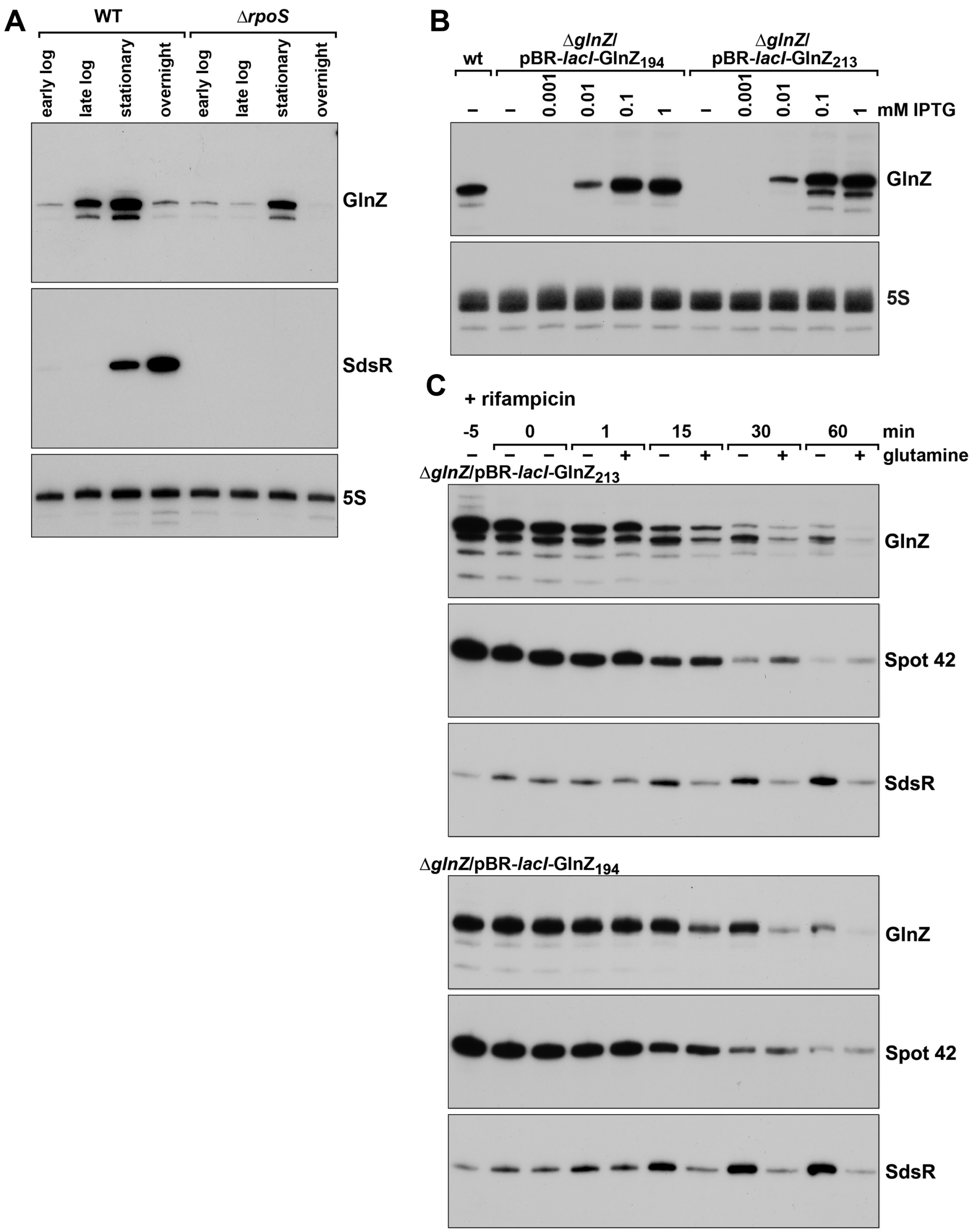

**Figure S2.** Effect of Δ*rpoS* and *glnZ* promoter on GlnZ levels. (**A**) The effect of Δ*rpoS* on GlnZ levels was evaluated by growing WT *crl*⁻ MG1655 and Δ*rpoS* (GSO108) strains to early logarithmic ($OD_{600}$~0.2), late logarithmic ($OD_{600}$~0.7), stationary phase ($OD_{600}$~3.5) or overnight ($OD_{600}$~3.5) in Gutnick medium with 0.4% glucose and 15 mM ammonium. Total RNA was collected and subjected to northern blot analysis as in Figure 1 with labeled oligonucleotide probes to GlnZ, SdsR and 5S. (**B**) WT MG1655 and Δ*glnZ* (GSO1153) cells harboring pBR-*lacI*-$GlnZ_{194}$ or pBR-*lacI*-$GlnZ_{213}$ were grown to stationary phase ($OD_{600}$ ~ 3.5) in Gutnick medium with 0.4% glucose and 15 mM ammonium and induced with a range of IPTG concentrations. Samples were taken one h post-induction. Total RNA was collected and subjected to northern blot analysis as in Figure 1 with labeled oligonucleotide probes to GlnZ and 5S. (**C**) The Δ*glnZ* mutant (GSO1153) strain harboring pBR-*lacI*-$GlnZ_{213}$ or pBR-*lacI*-$GlnZ_{194}$ was cultured to late logarithmic phase and expression from the plasmids was induced with 100 μM IPTG for 1 h. The cultures were treated with rifampicin, then split five min post rifampicin addition with 15 mM glutamine spiked into one culture. Cells were collected at the times indicated, and total RNA isolated was subjected to northern blot analysis as in Figure 1 with labeled oligonucleotide probes to GlnZ and 5S.

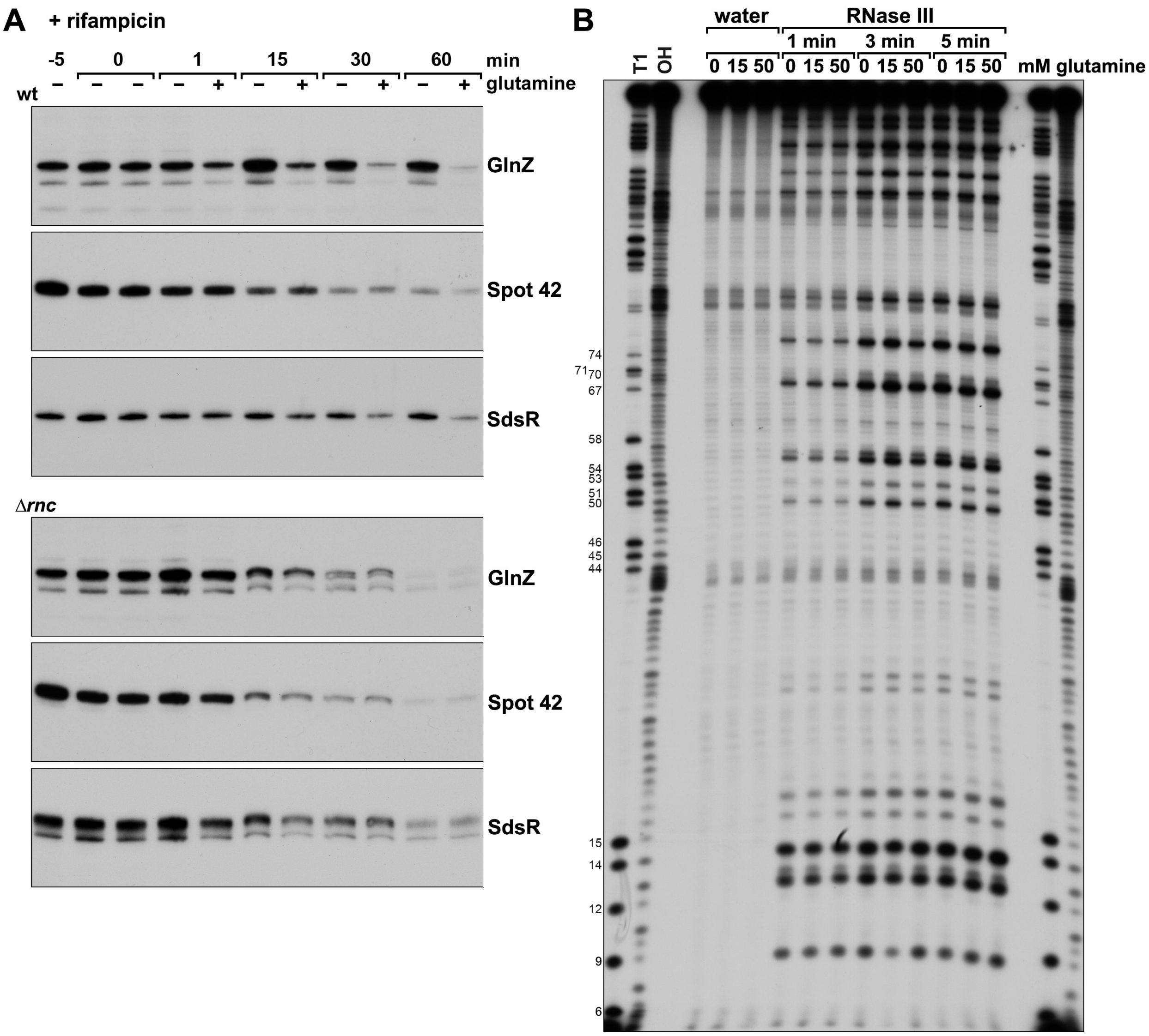

**Figure S3.** Effect of glutamine on RNase III cleavage in vivo and in vitro. (**A**) Wild type and Δ*rnc* strains (GSO1154) were cultured to stationary phase ($OD_{600}$~3.5) in Gutnick medium with 0.4% glucose and 15 mM ammonium and treated with rifampicin. Five min post rifampicin treatment, cultures were split, and 15 mM glutamine was spiked into one culture. Cells were collected at the times indicated and total RNA isolated was subjected to northern analysis. (**B**) RNase III in vitro cleavage assay was performed with $GlnZ_{194}$ transcribed in vitro from a T7 promoter and then radiolabeled. RNA was pre-incubated with either no glutamine, 15 mM or 50 mM glutamine for 1 h. Subsequently, water (as a control) or RNase III was added, and samples were incubated for 1, 3, or 5 min at 37˚C before stop solution was added. Samples were then resolved on a urea polyacrylamide gel, with T1 and OH ladders. Black dots denote strong cleavage by RNase III.

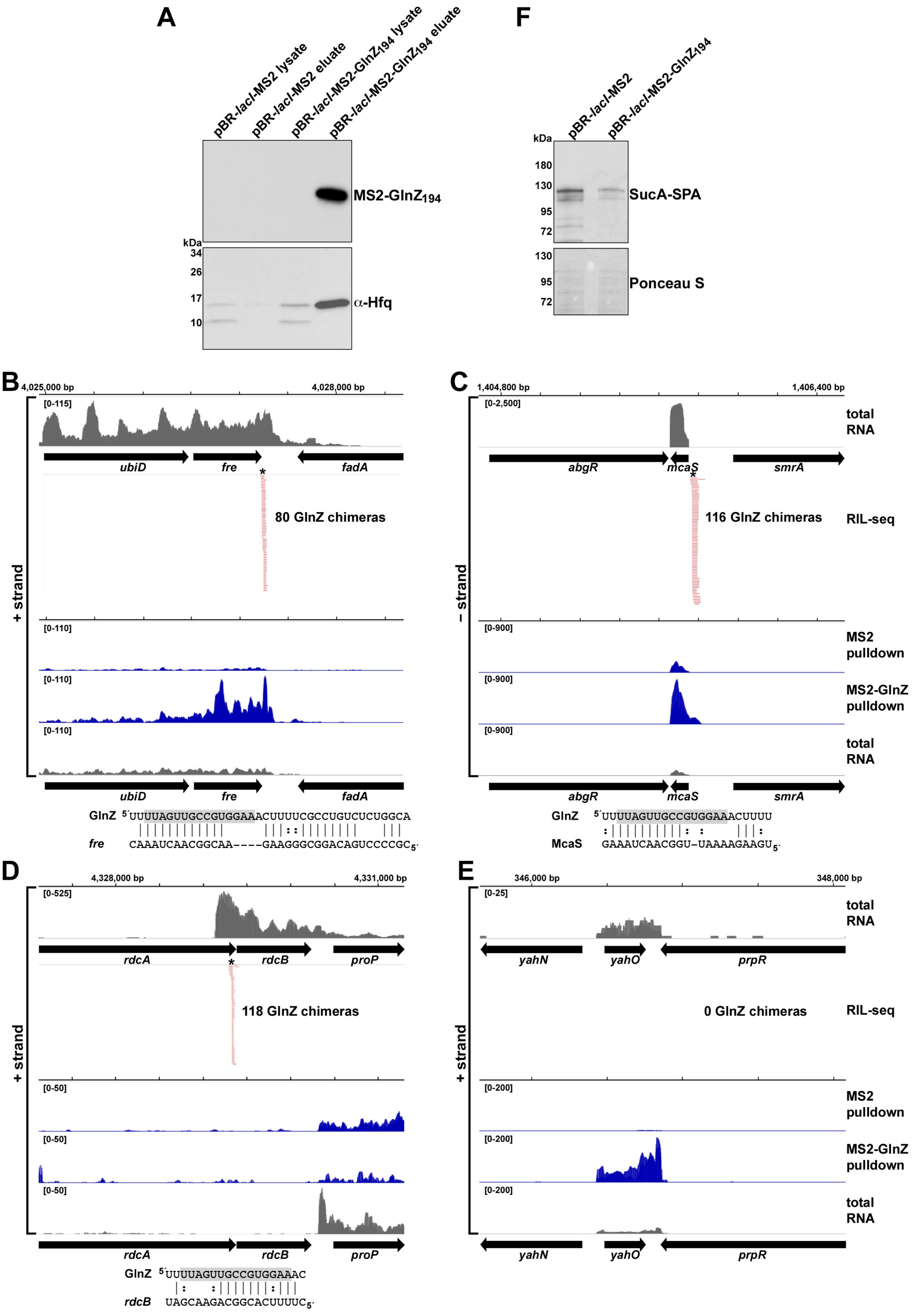
A
pBR-lacI-MS2 lysate
pBR-lacI-MS2 eluate
pBR-lacI-MS2-GlnZ194 lysate
pBR-lacI-MS2-GlnZ194 eluate
MS2-GlnZ194
kDa
34
26
17
10
α-Hfq
F
pBR-lacI-MS2
pBR-lacI-MS2-GlnZ194
kDa
180
130
95
72
SucA-SPA
130
95
72
Ponceau S
B
4,025,000 bp
4,028,000 bp
[0-115]
ubiD
fre
fadA
80 GlnZ chimeras
[0-110]
[0-110]
[0-110]
+ strand
ubiD
fre
fadA
GlnZ 5´UUUUAGUUGCCGUGGAAACUUUUCGCCUGUCUCUGGCA
fre CAAAUCAACGGCAA----GAAGGGCGGACAGUCCCCGC5´
C
1,404,800 bp
1,406,400 bp
[0-2,500]
total RNA
abgR
mcaS
smrA
116 GlnZ chimeras
RIL-seq
– strand
[0-900]
MS2 pulldown
[0-900]
MS2-GlnZ pulldown
[0-900]
total RNA
abgR
mcaS
smrA
GlnZ 5´UUUUAGUUGCCGUGGAAACUUUU
McaS GAAAUCAACGGU-UAAAAGAAGU5´
D
4,328,000 bp
4,331,000 bp
[0-525]
rdcA
rdcB
proP
118 GlnZ chimeras
+ strand
[0-50]
[0-50]
[0-50]
rdcA
rdcB
proP
GlnZ 5´UUUUAGUUGCCGUGGAAAC
rdcB UAGCAAGACGGCACUUUUC5´
E
346,000 bp
348,000 bp
[0-25]
total RNA
yahN
yahO
prpR
0 GlnZ chimeras
RIL-seq
+ strand
[0-200]
MS2 pulldown
[0-200]
MS2-GlnZ pulldown
[0-200]
total RNA
yahN
yahO
prpR

**Figure S4.** RNAs associated with $GlnZ_{194}$ identified by MAPS. (**A**) Northern blot and immunoblot assay after performing the MAPS co-purification protocol with lysates from Δ*glnZ* (GSO1153) overexpressing either MS2 or MS2-$GlnZ_{194}$. Lysate and eluate of indicated samples after 1 h overexpression with 1 mM IPTG were probed for GlnZ or Hfq. Browser images depict RIL-seq datasets from Melamed & Adams et al 2020 or MAPS data for GlnZ interactions with (**B**) *fre*, (**C**) McaS, (**D**) *rdcA,* and (**E**) *yahO*. Total RNA is depicted in grey, RIL-seq chimeras detected in M63 medium in red, and MAPS data for cultures grown in Gutnick medium with 0.4% glucose and 15 mM ammonium at stationary phase is depicted in blue for the pulldown with either MS2 or MS2-GlnZ overexpression. Predicted regions of interaction with GlnZ are depicted with the GlnZ seed sequence predicted by RIL-seq data highlighted in grey. The position of the predicted interaction is indicated by an asterisk in the browser images. (**F**) Immunoblot analysis of SucA-SPA levels. Δ*glnZ* (GSO1153) cells harboring pBR-*lacI*-MS2 or pBR-*lacI*-MS2-$GlnZ_{194}$ were grown to late logarithmic phase ($OD_{600}$ ~1.0) in Gutnick medium with 0.4% glucose and 15 mM ammonium and plasmids were induced with 1 mM IPTG. Samples were normalized by $OD_{600}$, separated by SDS-PAGE, and analyzed using immunoblot analysis. The Ponceau S-stained membrane serves as the loading control.

***glnA* 3′** (- strand)

GTACTACAGCGTCtaaGTGTTTTAGTTGCCGTGGAAACTTTTCGCCTGTCTCTGGCAGGCCTGGGATCGGTGGCAAG
CACATCACGCCGGATGCGACGCAAATGCGTCTTATCCGGCCTACACGGTGATGATGTGGTAGGCCGGAGCAGGTGAG
TCGCTCTCCAACGTGAAGTTTGTCAGCTATCTGTAGCCCATCTCTGCATGGGCTTTTTT

***glnP* 5′** (- strand)

taaTAACGCTACACCTGTAAAACGCACTGGCAGTTCCCTCTCCCCTATGGGGAGAGGATTAGGGTGAGGGGCGCAAA
CCCGCTCCGGGGCCATTAATTACCCTGAATTTGATTATTTACACCACGGTAACAGGAACAACATatgCAGTTTGACT
GGAGTGCCATCTGGCCTGCCATTCCGC

***sucA* 5′** (+ strand)

taaACCGTAGGCCTGATAAGACGCGCAAGCGTCGCATCAGGCAACCAGTGCCGGATGCGGCGTGAACGCCTTATCCG
GCCTACAAGTCATTACCCGTAGGCCTGATAAGCGCAGCGCATCAGGCGTAACAAAGAAATGCAGGAAATCTTTAAAA
ACTGCCCCTGACACTAAGACAGTTTTTAAAGGTTCCTTCGCGAGCCACTACGTAGACAAGAGCTCGCAAGTGAACCC
CGGCACGCACATCACTGTGCGTGGTAGTATCCACGGCGAAGTAAGCATAAAAAAGATGCTTAAGGGATCACGatgCA
GAACAGCGCTTTGAA

***aceE* 5′** (+ strand)

tagTGATTTTTCTGGTAAAAATTATCCAGAAGATGTTGTAAATCAAGCGCATATAAAAGCGCGGCAACTAAACGTAG
AACCTGTCTTATTGAGCTTTCCGGCGAGAGTTCAATGGGACAGGTTCCAGAAAACTCAACGTTATTAGATAGATAAG
GAATAACCCatgTCAGAACGTTTCCCAAATGACGTGGATCCGATCGAAA

***tmaR* (*yeeX*)** (- strand)

taaTTCCTGAACTTCAGAATCATCTTGCTGCTGCTTCGATTCAGCAAGGATAAAGGGTATGATAGTGAAAAGGGATA
AAAGCATTGTCATCTGCGGCAGCTATGAGTAATGTTGGCCCTAACGAATAGCGGTTGCTTAAACGAATCCGACTCTC
ACATTATCAGGGGTATAAAAatgGAAACTACCAAGCCTTCATTCCAGGACGTACTGGAATTTGTTCGTCTGTTCCGT
CGTAAGAACAAACTGCAACGTGAAATTCAGGACGTTGAGAAAAAGATCCGTGACAACCAGAAGCGCGTCCTGCTGCT
GGACAACCTGAGCGATTACATCAAGCCGGGGATGAGCGTTGAAGCAATCCAGGGCATCATCGCCAGCATGAAAGGTG
ACTATGAAGATCGCGTTGACGATTACATCATCAAAAATGCCGAGCTCTCCAAAGAACGCCGCGATATCTCCAAAAAG
CTGAAAGCTATGGGCGAAATGAAAAACGGCGAAGCGAAGtaaTTCCCGTTTTATTCAATGAGGGTTGCCCGGCAACC
CTCATTGCTCATTGA

***fre* 3′** (+ strand)

tgaTGCGCGTTTGTTTTGCCCTATTTATCGATCCGACAGAGAAAGCGCatgACAACCTTAAGCTGTAAAGTGACCTC
GGTAGAAGCTATCACGGATACCGTATATCGTGTCCGCATCGTGCCAGACGCGGCCTTTTCTTTTCGTGCTGGTCAGT
ATTTGATGGTAGTGATGGATGAGCGCGACAAACGTCCGTTCTCAATGGCTTCGACGCCGGATGAAAAAGGGTTTATC
GAGCTGCATATTGGCGCTTCTGAAATCAACCTTTACGCGAAAGCAGTCATGGACCGCATCCTCAAAGATCATCAAAT
CGTGGTCGACATTCCCCACGGAGAAGCGTGGCTGCGCGATGATGAAGAGCGTCCGATGATTTTGATTGCGGGCGGCA
CCGGGTTCTCTTATGCCCGCTCGATTTTGCTGACAGCGTTGGCGCGTAACCCAAACCGTGATATCACCATTTACTGG
GGCGGGCGTGAAGAGCAGCATCTGTATGATCTCTGCGAGCTTGAGGCGCTTTCGTTGAAGCATCCTGGTCTGCAAGT
GGTGCCGGTGGTTGAACAACCGGAAGCGGGCTGGCGTGGGCGTACTGGCACCGTGTTAACGGCGGTATTGCAGGATC
ACGGTACGCTGGCAGAGCATGATATCTATATTGCCGGACGTTTTGAGATGGCGAAAATTGCCCGCGATCTGTTTTGC
AGTGAGCGTAATGCGCGGGAAGATCGCCTGTTTGGCGATGCGTTTGCATTTATCtgaGATATAAAAAAACCCGCCCC
TGACAGGCGGGAAGAACGGCAACTAAACTGTTATTCAGTGGCATTTAGATCTATGACGTATCTGGCAAACCATGCCC
GATGCGACGCTGTCGCGTCTTATCGTGCCTACAAATAGTCCGAACCGTAGGCCGGATAAGGCGTTTACGCCGCATCC
GGCAATTGGTGCATGATGCCTGATGCGACGCTGTCGCGTCTTATCGTGCCTACAAATAGTCCGAACCGTAGGCCGGA
TAAGGCGTTTACGCCGCATCCGGCAATTGGTGCATGATGCCTGATGCGA

**McaS** (- strand)

GATATGATAACCAGACCGGGTCGGTCCAACAACGTATTACCCAAATTTCCAGTAATAAGTTCCAAATATTGCCGATA
TTTTAAGCAAAATACTTATGCATGATTATTCATTCACGATATTAATAATGTAACTTATATTTTCGTGAAATCTGTCA
CTGAAGAAAATTGGCAACTAAAGGTTAAAACCGTTATAACACAGTCaccGGCGCAGAGGAGACAATGCCGGATTTAA
GACGCGGATGCACTGCTGTGTGTACTGTAGAGTCTGGCGGATGCGGCAGTTCATCGACAGACTCTATTTTttt

***rdcB* (*yjcZ*) regulator of diguanylate cyclase** (+ strand)
TT**A**CGGCAGGCATGAATACTTTTTTCGCTGAGTTCGCTTCATGTTTGACGGAATTACAGACGCGTTTACGCGAAAGT
CTGGCTCTGCGTCAACAAAATGAATCGGTGGTCAGGCTGATGCAGCAGCAATTGCAGCAGACTGTGATGACTCACGG
CTGGATTTACACCGACGCCCAGCTGTTACGCGATGATATTCAAACACTTTTCACGGC*AGAA*CGATAttgACCAAGAC
GTTACTTGACGGCCCCGGTCGCGTGCTGGAGTCGGTTTATCCCCGCTTTTTAGTGGATCTGGCGCAGGGTGATGATG
CCCGCCTTCCACAAGCCCATCAGCAGCAGTTTCGTGAACGACTGATGCAGGAACTTCTTTCGCGTGTGCAGCTTCAG
ACATGGACGAACGGCGGCATGTTAAATGCGCCGCTTAGCCTGCGTCTGACATTGGTGGAAAAACTGGCGTCGATGCT
GGATCCCGGTCATCTGGCACTGACGCAGATCGCGCAGCATCTGGCGCTGCTGCAAAAAATGGATCACCGCCAGCACT
CTGCTTTCCCGGAGCTCCCCCAGCAAATTGCCGCCTTGTATGAGTGGTTTTCAGCCCGTTGTCGCTGGAAGGAAAAG
GCGTTAACGCAACGAGGCCTACTGGTGCAGGCAGGTGATCAGAGCGAGCAAATTTTTACCCGCTGGCGTGCTGGGGC
GTATAACGCCTGGTCGTTGCCTGGGCGCTGTTTTATCGTTCTGGAGGAGTTGCGCTGGGGGGCATTTGGCGATGCCT
GCCGTCTGGGAAGCCCGCAAGCGGTGGCGTTGTTGCTGGGTGATTTGCTCGAGAAAGCGACACAACATCTGGCAGAG
AGTATCAATGCGGCACCGACCACGCGTCACTATTACCATCAGTGGTTTGCCTCTTCGACCGTTCCGACGGGCGGGGA
GCATGCTGATTTTTTAAGTTGGCTGGGAAAGTGGACCACGGCAGATAAACAACCCGTTTGCTGGTCAGTGACCCAAC
GCTGGCAAACTGTCGCGCTGGGGATGCCACGACTCTGTTCAGCGCAGCGTCTGGCGGGGGCAATGCTCGAGGAAATC
TTCTCTGTAAATTTGGCGtaaATAATCAGTTACATCAATGAGTCCTAAACGAAATCCATGTGTGAAGTTGATCACAA
ATTTAAACACTGGTAGGGTAAAAAGGTCATTAACTGCCCAATTCAGGCGTCAACTG*G*TTTGATTGTACATTCCTTAA
CCGGAGGGTGTAAGCAAACCCGCTACGCTTGTTACAGAGATTGCATCCTGCAATTCCCGCTCCCCTT

***yahO* DUF1471 domain-containing protein** (+ strand)
ATGCAAAGGATCCATAGTGATTTCATCCATAAATAAGTGAACTAACTGCATCATATTTCTACCAAAAATAATCGGGT
GCGAGAGAGATCACAAAGTGTCTTATTTCCGGTTACTGGCGTTTATGCCCTGACTGAACTAATTATTAATC**A**ACCCA
ATAATGTGGGTGGGTGATAGTGTGATAACAACTCTGGAGCCGTAATatgAAAATAATCTCTAAAATGTTAGTCGGTG
CGTTAGCGTTAGCCGTTACCAATGTCTATGCCGCTGAATTGATGACCAAAGCGGAATTTGAAAAAGTTGAATCGCAG
TATGAAAAAATAGGTGATATTTCAACCAGCAATGAAATGTCGACTGCAGATGCAAAAGAAGATTTGATCAAAAAAGC
GGATGAAAAAGGGGCTGATGTGTTGGTACTGACCTCCGGTCAAACTGACAATAAGATCCACGGCACGGCAAATATTT
ATAAGAAGAAGtaaTTCTGAATCCTATGTAAACATCTCCGATGCGTAAGTTTATCGGTGATCATCTATTGAAATTTA
TGCCGGATAAAGCGTTCGCGCTGCATTCGGCAGTT*CA*GCTTTTCAGCCGCCGCCAGAACGTCGTCCGGCTGATGCCT
AAATAATTCGCCGCTGCTGTTTTATCGCCATTAAATTTCTCCAGTGCCTGTTGTGGTGTC

**Figure S5.** Sequences of *glnZ* as well as *sucA*, *glnP*, *aceE, tmaR, fre,* McaS, *rdcB*, and *yahO* near predicted regions of base pairing. Stop codons are indicated in red font, start codons are indicated in green font, and possible ribosome binding sites and stems defining terminators are in italics. Transcription start sites from (1) are indicated in bold. Underlined nucleotides indicate positions of RNase III cleavage from (2,3). Sequences highlighted in blue correspond to possible regions of pairing with the GlnZ seed sequence and sequences in orange font are ARN sequences that could be Hfq binding sites.

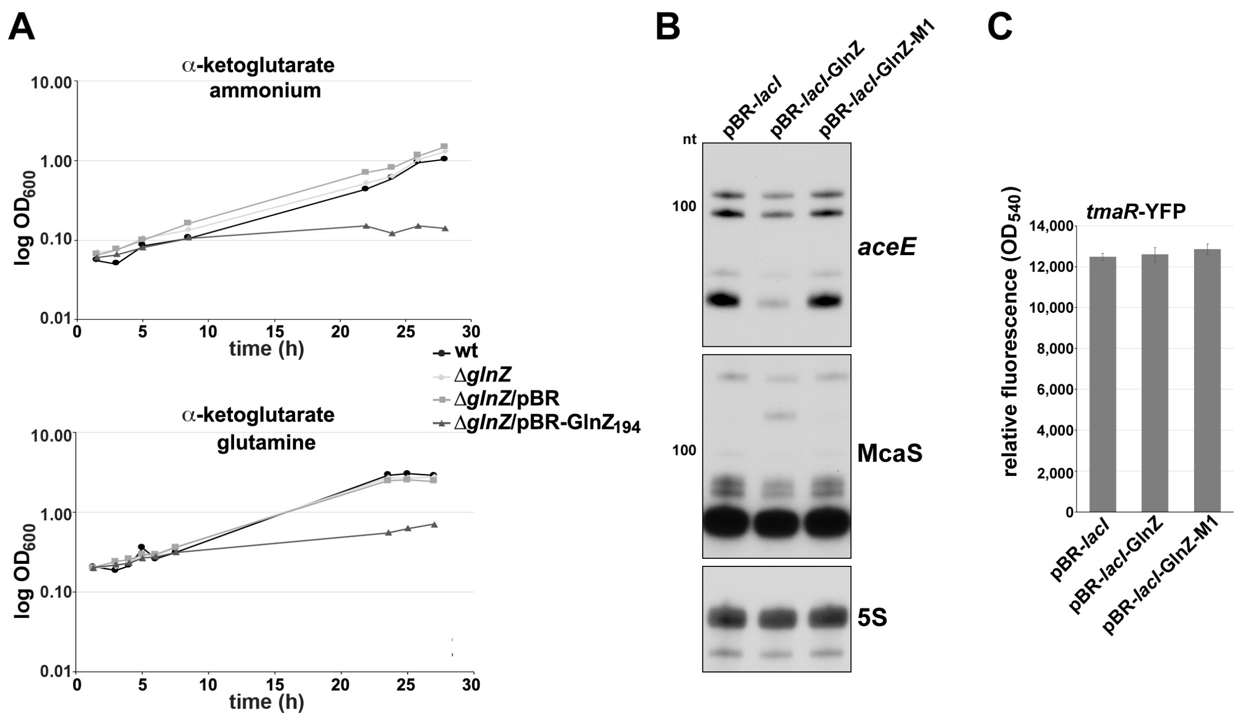

**Figure S6.** Growth assays and GlnZ regulation of *aceE* and McaS. (**A**) WT and Δ*glnZ* (GSO1153) strains harboring the plasmids indicated were subcultured to an $OD_{600}$ ~0.05 in Gutnick medium with 0.4% α-ketoglutarate as the carbon source and either 15 mM ammonium or 15 mM glutamine as the nitrogen source. Plasmids with induced with 1 mM IPTG. Cultures were incubated for 28 h and $OD_{600}$ was measured at the times indicated. The average of three independent replicates is plotted. (**B**) GlnZ downregulation of *aceE* and McaS transcripts. WT and Δ*glnZ* (GSO1153) strains harboring the indicated plasmids were grown in Gutnick medium with 0.4% glucose and 15 mM ammonium to stationary phase ($OD_{600}$ ~3.5). Plasmids were induced for 1 h with 1 mM IPTG. Total RNA was collected and subjected to northern blot analysis with labeled oligonucleotide probes to *aceE*, McaS, and 5S. For *aceE* and 5S, this figure shows the first three lanes of the northern presented in Figure 7D. The same blot was sequentially probed for McaS shown here. (**C**) $GlnZ_{194}$ does not regulate a *tmaR*-YFP fusion. The indicated strains were grown for 3 h in LB medium with 1 mM IPTG to induce WT and mutant GlnZ expression. Relative fluorescence units were determined by measuring $OD_{540}$ for each sample and normalizing by $OD_{600}$. The average of three independent replicates is shown with error bars indicating one standard deviation.

# A

### Class I

```
Providencia stuartii          GACCCCGCATCCATTAGAATTTGAAATGTACTACAGTGTGTAAC----------------
Yersinia pestis               GACTCCGCATCCTGTTGAGTTCGAACTGTATTACAGCGTTTAAGTTTACCTAATAGATTT
Shigella boydii               GACTCCGCATCCGGTAGAGTTTGAGCTGTACTACAGCGTCTAAT--AGTTGAAGTTGTAC
Escherichia coli O157:H7      GACTCCGCATCCGGTAGAGTTTGAGCTGTACTACAGCGTCTAAT--AGTTGAAGTTGTAC
Edwardsiella tarda            GACACCGCACCCGGTCGAGTTTGAACTCTACTACAGCGTATAACGCGACGCACGCCAGGC
Dickeya dadantii              GACGCCACATCCGGTTGAGTTCGAACTGTACTACAGCGTGTAATTCCCGCGCTTGGACGG
Erwinia tasmaniensis          GACGCCACACCCGGTTGAGTTCGAACTGTACTACAGCGTTTAAT----------TTTTGT
Cronobacter turicensis        GACGCCGCACCCGGTAGAGTTCGAACTCTACTACAGCGTGTAATAATTTTTTGATGTTGA
Sodalis glossinidius          AACGCCGCACCCGGTAGAGTTCGAGCTGTATTACAGCGTTTAAT----------------
Pantoea vagans                GACGCCACACCCGGTTGAGTTCGAGCTTTACTACAGCGTTTAAC---------------T
Proteus mirabilis             GGCACCACATCCACTTGAATTTGAAATGTATTACAGTGCGTAAC----------------
Xenorhabdus nematophila       GACACCACATCCTCTGGAATTTGAACTGTATTACAGTGTGTGAT----------------
Photorhabdus asymbiotica      GACACCACATCCACTGGAGTTTGAACTGTATTACAGTGTGTAAA----------------
                                * ** ** **  * ** ** **  * ** ***** *  * *

Providencia stuartii          -----------ACGCCGTTCCACTGCT------------------------------AGT
Yersinia pestis               CAAGGTGTAGGAAGGTGGCGAGCGAAAGCAGCCAACACACCGACAACTTGAAAGATGAAA
Shigella boydii               TACCCGGCGCAACAAC-------------------------------------GCCGGGA
Escherichia coli O157:H7      TACCCGGCGCAACAAC-------------------------------------GCCGGGA
Edwardsiella tarda            GACCCGGCGCCGTACCGGGTCCTTGCCG--------AACGCTCCACGCGAAGAGATCCGT
Dickeya dadantii              CAGGACGAAGAAAATAATGTTCACATTGCCGTCCGATAACCGTTTTCTGTTTATGTTTAT
Erwinia tasmaniensis          TATCCGGCGATATGTCATCTCGCAACTGTGGA------------ATGAGGTACGTCGGAT
Cronobacter turicensis        TAGCCAGCG-GATTTCAGGTTGTCGCCACTGCGTGAAATCCGC----------GAATGTT
Sodalis glossinidius          CACCCGA--------------------------------------------------GTT
Pantoea vagans                GGTCAGACG--------------------------------------------------T
Proteus mirabilis             ------GCACAATAGT------------------TAAAGTACTTACTTTGCTTATGATTT
Xenorhabdus nematophila       -GCCTGATCTTGTGCTGAATAGTTGTCCTTGATGATA-------GCTCTGTTATTCCTTT
Photorhabdus asymbiotica      TTAGCAGCGCCATTCCTGCT---------------------------------GATCATT

Providencia stuartii          CTTTGTTGCCGTG-AAAC-ATTTGCCCCATCTCTGATGGGGCCTTTTCTCC
Yersinia pestis               GGTTGTTGCCGTGGAAAC-TTTTGGCCCATCTTAGGATGGGCTTTTTTCTC
Shigella boydii               TTTAGTTGCCGTGGAAAC-TTTCAGCCCATCTCTGCATGGGCTTTTTTCTC
Escherichia coli O157:H7      TTTAGTTGCCGTGGAAAC-TTTCAGCCCATCTCTGCATGGGCTTTTTTCTC
Edwardsiella tarda            TTTTGTTGCCGTGGAAAC-TTTCAGCCCATCTTCGGATGGGTTTTTTTCGC
Dickeya dadantii              TTTTGTTGCCGTGGAAACTTTTCAGCCCATCTTCGGATGGGCTTTTTTCTC
Erwinia tasmaniensis          ATTTGTTGCCGTGGAAAC-TTTCAGCCCATCTTCGGATGGGCTTTTTTCTC
Cronobacter turicensis        TTTTGTTGCCGTGGAAAC--TTTAGCCCATCTTCGGATGGGCTTTTTTCAC
Sodalis glossinidius          TTTTGTTGCCGTGGAAAC-TTTCAGCCCATCTTCGGATGGGCTTTTTTCTC
Pantoea vagans                TTTTGTTGCCGTGGAAAC-TTTTAGCCCATCTTCGGATGGGCTTTTTTCTC
Proteus mirabilis             TTTTGTTGCCGTG-AAGCTATTTTGCCCATCTTCGGATGGGCTCTTTTCTC
Xenorhabdus nematophila       TTTTGTTGCCGTA-AAAC-TTTTGGCCCATCCTCGGATGGGTTCTTTTACC
Photorhabdus asymbiotica      TTTTGTTGCCGTGAAACC--TTTAGCCCATCCCCGGATGGGCCCTTTTCCC
                                * ********* ** *  **  *******   *   ***   ***   *
```

### Class II

```
Cedecea neteri                   GGTAGAGTTCGAGCTGTACTACAGCGTCTAAT----ATTTA------------GAAATCC
Klebsiella michiganensis         GGTAGAGTTCGAGCTGTACTACAGCGTCTAAT---AATTTA------------GAAATCC
Salmonella enterica              GGTAGAGTTTGAGCTGTACTACAGCGTTTAATCGTATATTA------------AAAATCC
Enterobacteriaceae bacterium     GGTAGAGTTCGAGCTGTACTACAGCGTTTAAT---AATTTA--------------AATCC
Phytobacter diazotrophicus       GGTAGAGTTCGAGCTGTACTACAGCGTTTAAT---AATTTA--------------AATCC
Enterobacter mori                AGTTGAATTCGAACTGTACTACAGCGTTTAAGTCAGAATTAAAGAAAGAAATTAATACTC
Kosakonia sp.                    GGTAGAGTTCGAACTGTACTACAGCGTTTAAT---AAATTA------------AATACTC
                                  ** ** ** ** ************** ***       ***            *  *

Cedecea neteri                   AACGAATTTCGCGTTGCAGCACGACGGCAACTGAGTGAATCCCTGGAAGCATAGGTAACT
Klebsiella michiganensis         AACGAATTTCGCGTTGCAGCAAGACGGCAACTGAGTGAATCCCTGGGAGCATAGAGAACT
Salmonella enterica              GACGAATTTCGCGTTGCTGCAAGGCGGCAACTGAGCACATCCCCGGGAGCATAGATAGCG
Enterobacteriaceae bacterium     GAGGGATTTCTTGTTACAGCAAGGCGGCGATTGAGTGAATCCCCGGGAGCATAGAGATCT
Phytobacter diazotrophicus       GAGGGATTTCTTGTTACAGCAAGGCGGCGATTGAGTGAATCCCCGGGAGCATAGAGATCT
Enterobacter mori                GGCGGATTTCTTGTCGCAGCAAGGCGGCAACTGAGTGAATCCCCGGGAGCATAGATAGCT
Kosakonia sp.                     GGCGGATTTCTTGTCGTAGCAAGACGGCAACTGAGCGAATCCCCGGGAGCATAGATAACT
                                    * *****  **     *** * **** * ****    ***** ** *******  * *

Cedecea neteri                   ATGTGACCAGGGTGAGTAAGGGCAGCCAACGCAGCTGTGGCGCGAAGGGCGTTAGGAG-T
Klebsiella michiganensis         ATGTGACCAGGGTGAATGAGGGCAGCCAACGTAGCGGTGGCGTGAAGGGCGTTAGGAG--
Salmonella enterica              ATGTGACCGGGGTAAGCGAAGGCAGCCAACGCAGCAGCAGCGTGAAAGGCGTCAGGAGTT
Enterobacteriaceae bacterium     ATGTGACCGGGGTAAGCGAAGGAAGCCAACGCCGCTGTAGCGAGAAAGACAT---AGGAT
Phytobacter diazotrophicus       ATGTGACCGGGGTAAGCGAAGGAAGCCAACGCCGCTGTAGCGAGAAAGACAT---AGGAT
Enterobacter mori                ATGTGACCGGGGTGAACGAAGGCAGCCAACGCGGCTGCGGCGAGAAAGACGCGAGAGGAT
Kosakonia sp.                     ATGTGACCGGGGTGAGCGAAGGCAGCCAACGCAGCAACGGCGAGAAAGACGCCAGAGGA-
                                 ******** **** *   * ** ********  **     *** *** * *        *

Cedecea neteri                   TTTTTGTTGCCGTGGAAACTTT-GGCCCATCTCCGGATGGGCTTTTTTCTCCACCGGAA
Klebsiella michiganensis         TTTTTGTTGCCGTGGAAACTTT-AGCCCATCTTCGGATGGGCTTTTTTCTCCATCGGA-
Salmonella enterica              TTTGAGTTGCCGTGGAAACTTTCAGCCCATCCCAGGATGGGCTTTTTTCTCCACCAACA
Enterobacteriaceae bacterium     ATTTTGTTGCCGTGGAAACTTTCAGCCCATCCT-AGATGGGCTTTTTTCTCCACCAACA
Phytobacter diazotrophicus       ATTTTGTTGCCGTGGAAACTTTCAGCCCATCCT-AGATGGGCTTTTTTCTCCACCAACA
Enterobacter mori                TTTTTGTTGCCGTGGAAACTTTCAGCCCATCTTCGGATGGGCTTTTTTCTCCACCAACA
Kosakonia sp.                    --TTTGTTGCCGTGGAA-CTTTTAGCCCATCCCTGGATGGGCTTTTTTCTCCACCAACA
                                   *  *********** ****   *******     ***************** *
```

### Class III

```
Escherichia coli K-12 MG1655     TAAGTGTTTTAGTTGCCGTGGAAACTTTTCGCCTGTCTCTGGCAGGCCTGGGATCGGTGG
Shigella_sp. PAMC 28760          TAAGTGTTTTAGTTGCCGTGGAAACTTTTCGCCTGTCTCTGGCAGGCCTGGGATCGGTGG
Salmonella sp. S13               TAAGTGTTTTAGTTGCCGTGGAAACTTTTCGCCTGTCTCTGGCAGGCCTGGGATCGGTGG
Salmonella sp. HNK130            TAAGTGTTTTAGTTGCCGTGGAAACTTTTCGCCTGTCTCTGGCAGGCCTGGGATCGGTGG
E. albertii 05-3106              TAAGTGTTTTAGTTGCCGTGGAAACTTTTCGCCTGTCCCTGGCAGGCTTGAGATCGATGG
E. albertii CB9786               TAAGTGTTTTAGTTGCCGTGGAAACTTTTCGCCTGTCCCTGGCAGGCTTGAGATCGATGG
E. albertii NIAH Bird 3          TAAGTGTTTTAGTTGCCGTGGAAACTTTTCGCCTGTCCCTGGCAGGCTTGAGATCGATGG
E. albertii Sample 166           TAAGTGTTTTAGTTGCCGTGGAAACTTTTCGCCTGTCCCTGGCAGGCTTGAGATCGATGG
                                 ****************************************** ********* ** ***** ***

Escherichia coli K-12 MG1655     CAAGCACATCACGCCGGATGCGACGCAAA-TGCGTCTTATCCGGCCTACACGGTGATGAT
Shigella_sp. PAMC 28760          CAAGCACATCACGCCGGATGCGACGCAAA-TGCGTCTTATCCGGCCTACACGGTGATGAT
Salmonella sp. S13               CAAGCACATCACGCCGGATGCGACGCAAA-TGCGTCTTATCCGGCCTACACGGTGATGAT
Salmonella sp. HNK130            CAAGCACATCACGCCGGATGCGACGCAAA-TGCGTCTTATCCGGCCTACACGGTGATGAT
E. albertii 05-3106              CAAGCACGTCATGCCGGATACGACGCAAATTGCATCTTATCCGGCCTACACGGTGATTAT
E. albertii CB9786               CAAGCACGTCATGCCGGATACGACGCAAATTGCATCTTATCCGGCCTACACAGTGATTAT
E. albertii NIAH Bird 3          CAAGCACGTCATGCCGGATACGACGCAAATTGCATCTTATCCGGCCTACACAGTGATTAT
E. albertii Sample 166           CAAGCACGTCATGCCGGATACGACGCAAATTGCATCTTATCCGGCCTACACAGTGATTAT
                                 ******* *** ******* ********* *** ***************** ***** **

Escherichia coli K-12 MG1655     GTGGTAGGCCGGAGCAGGTG-AGTCGCTCTCCAACGTGAAGTTTGTCAGCTATCTGTAGC
Shigella sp. PAMC 28760          GTGGTAGGCCGGAGCAGGTG-AGTCGCTCTCCAACGTGAAGTTTGTCAGCTATCTGTAGC
Salmonella sp. S13               GTGGTAGGCCGGAGCAGGTG-AGTCGCTCTCCAACGTGAAGTTTGTCAGCTATCTGTAGC
Salmonella sp. HNK130            GTGGTAGGCCGGAGCAGGTG-AGTCGCTCTCCAACGTGAAGTTTGTCAGCTATCTGTAGC
E. albertii 05-3106              GCGGTAGGCCGGAGCAGGTGAAGTCGCTCTCCGAAGTGAAGTTTGTCAGCTATCTGTAGC
E. albertii_CB9786               GCGGTAGGCCGGAGCAGGTGAAGTCGCTCTCCGAAGTGAAGTTTGTCAGCTATCTGTAGC
E. albertii NIAH Bird 3          GCGGTAGGCCGGAGCAGGTGAAGTCGCTCTCCGAAGTGAAGTTTGTCAGCTATCTGTAGC
E. albertii Sample 166           GCGGTAGGCCGGAGCAGGTGAAGTCGCTCTCCGAAGTGAAGTTTGTCAGCTATCTGTAGC
                                 * ****************** *********** * *************************

Escherichia coli K-12 MG1655     CCATCTCTGCATGGGCTTTTTT
Shigella sp. PAMC 28760          CCATCTCTGCATGGGCTTTTTT
Salmonella sp. S13               CCATCTCTGCATGGGCTTTTTT
Salmonella sp. HNK130            CCATCTCTGCATGGGCTTTTTT
E. albertii 05-3106              CCATCTCTGCATGGGCTTTTTT
E. albertii_CB9786               CCATCTCTGCATGGGCTTTTTT
E. albertii NIAH Bird 3          CCATCTCTGCATGGGCTTTTTT
E. albertii Sample 166           CCATCTCTGCATGGGCTTTTTT
                                 **********************
```

# B

```
Proteus mirabilis                 TTTTGTTGCCGTG-AAGC
Providencia stuartii              CTTTGTTGCCGTG-AAAC
Xenorhabdus nematophila           TTTTGTTGCCGTA-AAAC
Photorhabdus asymbiotica          TTTTGTTGCCGTGAAACC
Yersinia pestis                   GGTTGTTGCCGTGGAAAC
Salmonella enterica               TTGAGTTGCCGTGGAAAC
Shigella boydii                   TTTAGTTGCCGTGGAAAC
Escherichia coli O157:H7          TTTAGTTGCCGTGGAAAC
Escherichia coli K-12 MG1655      TTTAGTTGCCGTGGAAAC
Shigella sp. PAMC 28760           TTTAGTTGCCGTGGAAAC
Salmonella sp.                    TTTAGTTGCCGTGGAAAC
Kosakonia sp.                     TTTTGTTGCCGTGGAA-C
Edwardsiella tarda                TTTTGTTGCCGTGGAAAC
Dickeya dadantii                  TTTTGTTGCCGTGGAAAC
Cronobacter turicensis            TTTTGTTGCCGTGGAAAC
Sodalis glossinidius              TTTTGTTGCCGTGGAAAC
Pantoea vagans                    TTTTGTTGCCGTGGAAAC
Cedecea neteri                    TTTTGTTGCCGTGGAAAC
Klebsiella michiganensis          TTTTGTTGCCGTGGAAAC
Enterobacteriaceae bacterium      TTTTGTTGCCGTGGAAAC
Phytobacter diazotrophicus        TTTTGTTGCCGTGGAAAC
Enterobacter mori                 TTTTGTTGCCGTGGAAAC
Erwinia tasmaniensis              ATTTGTTGCCGTGGAAAC
                                      ********  ** *
```

**Figure S7.** Conservation of GlnZ seed sequence and terminator despite overall lack of conservation. (**A**) Sequence alignment of the 3´UTR of *glnA* from various Gammaproteobacteria containing class I, class II, or class III GlnZ homologs. The *glnA* stop codon is indicated with a box. Nucleotides conserved in all sequences are indicated with an asterisk. The GlnZ seed sequence predicted by RIL-seq data is highlighted in grey, and the seed residues conserved across all species used in our analysis are in red font. The predicted terminator stem-loops are in orange font. (**B**) Sequence alignment of seed sequence from Gammaproteobacteria. All alignments were generated using MUSCLE (4).

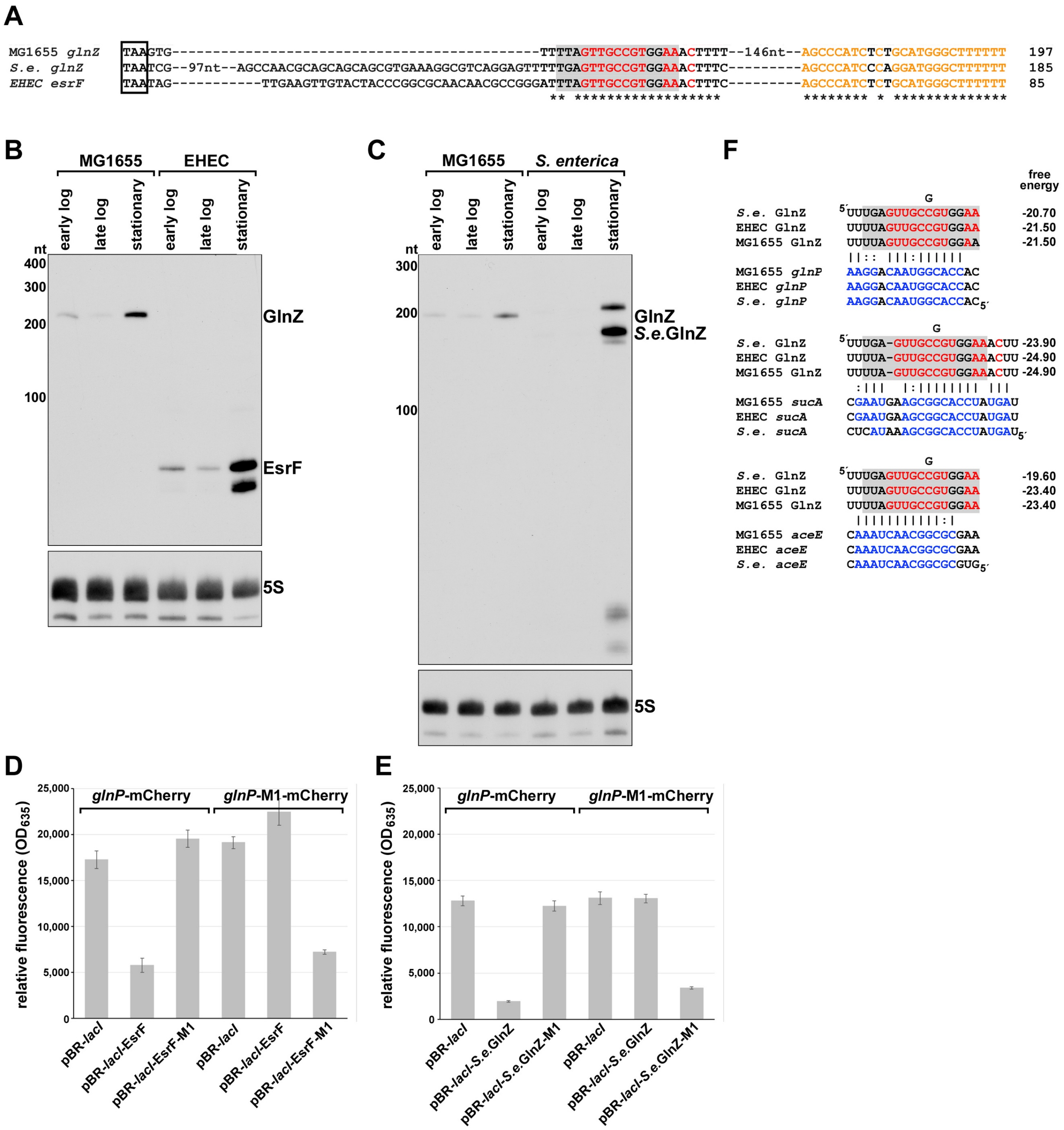

**Figure S8.** Conservation of GlnZ expression, and function. (**A**) Sequence alignment of the GlnZ seed sequence region (seed sequence predicted by RIL-seq data highlighted in grey, and residues conserved across all species used in our analysis in red font) and terminator stem-loop (conserved residues in orange font) for the 3′ UTRs of *glnA* from *E. coli* MG1655, EHEC (NC_002695.2) and *Salmonella enterica* (NC_003197.2). Nucleotides conserved in all three sequences are indicated with an asterisk. Positions are given relative to the *glnA* stop codon. Alignment was generated using MUSCLE (4). (**B**) Northern blot analysis of GlnZ levels with *E. coli* MG1655 or EHEC cultured to early logarithmic, late logarithmic, or stationary phases in Gutnick medium with 0.4% glucose and 15 mM ammonium. RNA was isolated from samples and subjected to northern analysis as described in Figure 1. (**C**) Northern blot analysis of GlnZ levels with *E. coli* MG1655 or *S. enterica* LT2 cultured to early logarithmic, late logarithmic, or stationary phases in Gutnick medium with 0.4% glucose and 15 mM ammonium. RNA was isolated from samples and subjected to northern analysis as described in Figure 1. Regulation by (**D**) EsrF-M1 and (**E**) *S.e.*GlnZ-M1 is restored for the *glnP-M1-mCherry* fusion. The indicated strains were grown for 3 h in Gutnick medium with 0.4% glucose and 15 mM ammonium and 1 mM IPTG to induce WT and mutant EsrF and *S.e.*GlnZ expression and either 0.2% arabinose (*glnP-mCherry*) or 2% arabinose (*glnP-M1-mCherry*) to induce the mCherry fusion. Relative fluorescence units were determined by measuring $OD_{635}$ for each sample and normalizing by $OD_{600}$. The average of three independent replicates are plotted. Error bars indicate one standard deviation. (**F**) Conservation of the region of base pairing with GlnZ targets *glnP*, *sucA*, and *aceE*. The sequences of the base pairing regions for the targets are shown with the GlnZ seed sequence predicted by RIL-seq data highlighted in grey, the GlnZ residues conserved across all species used in our analysis in red font, and the conserved target nucleotides involved in base pairing in blue font. The M1 mutant sequence is indicated above the alignments. The interaction between GlnZ and its targets in *E. coli* is indicated with lines.

### Supplementary Tables

**Table S1.** Strains used in study

| GSO Name | Strain Name | Relevent Genotype | Source |
|---|---|---|---|
| GSO983 | MG1655 | *Escherichia coli* wild-type strain (*crl-*) | Lab stock |
| GSO982 | MG1655 | *Escherichia coli wild-type strain* (*crl+*) | Lab stock |
| | NM400 | MG1655 mini-λ $cm^R$ ts | Gottesman Lab |
| | NM500 | MG1655 mini-λ $tet^R$ ts | Gottesman Lab |
| | PM1205 | PM1203 *lacI'*:: $P_{BAD}$*-cat-sacB:lacZ,* miniλ $tet^R$ | (5) |
| | NRD1166 | MG1655 Zeo T1 T2 $P_{lac}$-*ccdB-kan*-mCherry at *lac* locus, mini-λ:tet | Gottesman Lab |
| GSO108 | MG1655 Δ*rpoS* | MG1655 Δ*rpoS*::*tet* | (6) |
| | JW3839 Δ*glnG* | Δ*glnG::kan* (=Δ*ntrC::kan*) | Keio collection |
| GSO1151 | MG1655 Δ*glnG* | Δ*glnG::kan* | This study |
| GSO1152 | MG1655 Δ$P_{glnA}$ | Δ$P_{glnA}$*::kan* | This study |
| GSO1153 | MG1655 Δ*glnZ* | Δ*glnZ::kan* | This study |
| GSO1154 | MG1655 Δ*rnc* | Δ*rnc::cam* | Lab stock |
| GSO1155 | MG1655 Δ*rnc*Δ*glnZ* | Δ*rnc::cam* Δ*glnZ::kan* | This study |
| GSO1156 | PM1205 $P_{glnA}$*-lacZ* | $P_{glnA}$*:lacZ* | This study |
| GSO1157 | PM1205 $P_{glnZ}$*-lacZ* | $P_{glnZ}$*:lacZ* | This study |
| GSO1158 | NRD1166 *glnP*-mCherry | *glnP*-mCherry | This study |
| GSO1159 | NRD1166 *glnP*-M1-mCherry | *glnP*-M1-mCherry | This study |
| | DY330 *sucA-SPA* | W3110 Δ*lacU169 gal490 λCI857* Δ*(cro-bioA) sucA:SPA::kan* | (7) |
| GSO1160 | NM500 Δ*sucA::cat-sacB* | Δ*sucA::cat-sacB* | This study |
| GSO1161 | MG1655 *sucA*-SPA | *sucA-SPA::kan* | This study |
| GSO1162 | MG1655 *sucA*-M1-SPA | *sucA-M1-SPA::kan* | This study |
| GSO1163 | MG1655 Δ*rnc sucA*-SPA | Δ*rnc::cam sucA-SPA::kan* | This study |
| | SX1989 *yeeX-YFP* | *yeeX793-YFP::cat* | (8) |
| GSO1164 | MG1655 *tmaR-YFP* | *yeeX793-YFP::cat* | This study |
| | *Escherichia coli* EHEC O157:H7 EDL933 | | ATCC |
| | *Salmonella enterica* subs. e*nterica* LT2 | | Lab stock |
| GSO1165 | NEB5α + pBR-lacI-$GlnZ_{213}$ | pBR-lacI-$GlnZ_{213}$ | This study |
| GSO1166 | NEB5α + pBR-lacI-$GlnZ_{194}$ | pBR-lacI-$GlnZ_{194}$ | This study |
| GSO1167 | NEB5α + pBR-lacI-$GlnZ_{194}$-M1 | pBR-lacI-$GlnZ_{194}$-M1 | This study |
| GSO1168 | NEB5α + pBR-lacI-$GlnZ_{174}$ | pBR-lacI-$GlnZ_{174}$ | This study |
| GSO1169 | NEB5α + pBR-lacI-MS2-$GlnZ_{194}$ | pBR-lacI-MS2-$GlnZ_{194}$ | This study |
| GSO1170 | NEB5α + pBR-lacI-EsrF | pBR-lacI-EsrF | This study |
| GSO1171 | NEB5α + pBR-lacI-EsrF-M1 | pBR-lacI-EsrF-M1 | This study |
| GSO1172 | NEB5α + pBR-lacI-S.e.GlnZ | pBR-lacI-S.e.GlnZ | This study |
| GSO1173 | NEB5α + pBR-lacI-S.e.GlnZ-M1 | pBR-lacI-S.e.GlnZ-M1 | This study |
| GSO1174 | NEB5α + pBR-$GlnZ_{194}$ | pBR-$GlnZ_{194}$ | This study |

| | | | |
|---|---|---|---|
| GSO1175 | NEB5α + pBR-GlnZ$_{194}$-M1 | pBR-GlnZ$_{194}$-M1 | This study |
| GSO1176 | NEB5α + pUC19-SucA-SPA | pUC19-SucA-SPA | This study |
| GSO1177 | NEB5α + pUC19-SucA-M1-SPA | pUC19-SucA-M1-SPA | This study |

**Table S2.** Plasmids used in study

| **Plasmid Name** | **Plasmid Description** | **Source** |
|---|---|---|
| pBR | pBR322 carrying an inducible PlacO-1 promoter. AmpR KanR | (9) |
| pBR-*lacI* | pNM46, pBR carrying the *lacI* gene ($amp^R$) | N. Majdalani |
| pBR-*lacI*-$GlnZ_{213}$ | pBR-*lacI* carrying 213 nt variant of GlnZ | This study |
| pBR-*lacI*-$GlnZ_{194}$ | pBR-*lacI* carrying 194 nt variant of GlnZ | This study |
| pBR-*lacI*-$GlnZ_{194}$-M1 | pBR-*lacI* carrying 194 nt variant of GlnZ with M1 mutation | This study |
| pBR-*lacI*-$GlnZ_{174}$ | pBR-*lacI* carrying 174 nt variant of GlnZ | This study |
| pBR-*lacI*-MS2 | pBR-*lacI* carrying the MS2 tag | N. Thongdee |
| pBR-*lacI*-MS2-$GlnZ_{194}$ | pBR-*lacI* carrying 194 nt variant of GlnZ | This study |
| pBR-*lacI*-EsrF | pBR-*lacI* carrying EsrF | This study |
| pBR-*lacI*-EsrF-M1 | pBR-*lacI* carrying EsrF with seed sequence mutation | This study |
| pBR-*lacI*-*S.e.*GlnZ | pBR-*lacI* carrying *S. enterica* GlnZ | This study |
| pBR-*lacI*-*S.e.*GlnZ-M1 | pBR-*lacI* carrying *S. enterica* GlnZ with M1 mutation | This study |
| pBR-$GlnZ_{194}$ | pBR carrying 194 nt variant of GlnZ | This study |
| pBR-$GlnZ_{194}$-M1 | pBR carrying 194 nt variant of GlnZ with M1 mutation | This study |
| pBR-RyhB | pBR carrying sRNA *ryhB* | (10) |
| pUC19 | $amp^R$ | New England Biolabs |
| pUC19-SucA-SPA | pUC19 carrying *sucA-SPA::kan* including the 5′ UTR of SucA | This study |
| pUC19-SucA-M1-SPA | pUC19 carrying *sucA-SPA::kan* including the 5′ UTR of SucA with M1 mutation | This study |

**Table S3.** Oligonucleotides used in study

| Name | Purpose | Sequence |
|---|---|---|
| AK280 | Northern probe against GlnZ | ATGGGCTACAGATAGCTGACAAACTTCACG |
| LW030 | Northern probe against 5S rRNA | CGGCGCTACGGCGTTTCACTTCTG |
| MR023 | Northern probe against Spot 42 | GGTCTGAAAGATAGAACATCTTACCTCTGT |
| LW066 | Northern probe against SdsR | GTATTCGGTCCAGGGAAATGGCTCTTGGG |
| LW055 | Oligo for AceE primer extension analysis | GATCGGATCCACGTCATTTGGG |
| LW138 | Northern probe against EsrF and *S.e.*GlnZ | GCTGAAAGTTTCCACGGCAACTAAATCCCG |
| LW178 | Northern probe against AceE 5'UTR | CTCTCGCCGGAAAGCTCAATAAGACAGGTTCTACGTTTAGTTGCCGCGC |
| LW251 | Northern probe against McaS | TCCGCGTCTTAAATCCGGCATTGTCTCCTCTGCGCCGGT |
| LW043 | Forward primer to amplify pBR-*lacI* plasmid for Gibson cloning | GAATTCTCATGTTTGACAG |
| LW044 | Reverse primer to amplify pBR-*lacI* plasmid for Gibson cloning | GACGTCAGTATCTTGTTATC |
| LW045 | Forward primer to amplify $GlnZ_{213}$ for Gibson cloning into pBR-*lacI* | GATAACAAGATACTGACGTCGTACTACAGCGTCTAAGTG |
| LW046 | Reverse primer to amplify $GlnZ_{174}$/$GlnZ_{194}$/$GlnZ_{213}$ for Gibson cloning into pBR-*lacI* | GCTGTCAAACATGAGAATTCAAAAAAGCCCATGCAGAG |
| LW047 | Forward primer to amplify $GlnZ_{194}$ for Gibson cloning into pBR-*lacI* | GATAACAAGATACTGACGTCTTTTAGTTGCCGTGGAAAC |
| LW060 | Forward primer to amplify $GlnZ_{174}$ for Gibson cloning into pBR-*lacI* | GATAACAAGATACTGACGTCTTTCGCCTGTCTCTGGCAG |
| LW127 | Forward QuikChange primer to make seed sequence mutant in pBR-*lacI*-$GlnZ_{194}$ | AGGCGAAAAGTTTCCACCGCAACTAAAAGACGTCA |
| LW128 | Reverse QuikChange primer to make seed sequence mutant in pBR-*lacI*-$GlnZ_{194}$ | TGACGTCTTTTAGTTGCGGTGGAAACTTTTCGCCT |
| LW113 | Forward primer to amplify pBR-*lacI*-MS2 plasmid for Gibson cloning | AACCATTATTATCATGACATTAACCTATAAAAATAG |
| LW114 | Reverse primer to amplify pBR-*lacI*-MS2 plasmid for Gibson cloning | CAGACCCTGATGGTGTCTG |
| LW115 | Forward primer to amplify $GlnZ_{194}$ for Gibson cloning into pBR-lacI-MS2 | GCAGACACCATCAGGGTCTGTTTTAGTTGCCGTGGAAAC |
| LW116 | Reverse primer to amplify $GlnZ_{194}$ for Gibson cloning into pBR-lacI-MS2 | ATGTCATGATAATAATGGTTAAAAAAGCCCATGCAGAG |
| LW132 | Forward primer to amplify EsrF for Gibson cloning into pBR-*lacI* | GATAACAAGATACTGACGTCTAGTTGAAGTTGTACTACCC |
| LW133 | Reverse primer to amplify EsrF for Gibson cloning into pBR-*lacI* | GCTGTCAAACATGAGAATTCAAAAAAGCCCATGCAGAG |
| LW139 | Forward QuikChange primer to make seed sequence mutant in pBR-*lacI*-EsrF | GGCTGAAAGTTTCCACCGCAACTAAATCCCGGC |
| LW140 | Reverse QuikChange primer to make seed sequence mutant in pBR-*lacI*-EsrF | GCCGGGATTTAGTTGCGGTGGAAACTTTCAGCC |
| LW147 | Forward primer to amplify *S.e.*GlnZ for Gibson cloning into pBR-*lacI* | GATAACAAGATACTGACGTCTCGTATATTAAAAATCCGAC |

| | | |
|---|---|---|
| LW148 | Reverse primer to amplify *S.e.*GlnZ for Gibson cloning into pBR-*lacI* | GCTGTCAAACATGAGAATTCAAAAAAGCCCATCC |
| AK305 | Forward primer to clone $GlnZ_{213}$ into pBR | GACGTCGTACTACAGCGTCTAAGTGTTTTAGTTG |
| AK306 | Forward primer to clone $GlnZ_{194}$ into pBR | GACGTCTTTTAGTTGCCGTGGAAACTTTTCG |
| AK307 | Forward primer to clone $GlnZ_{174}$ into pBR | GACGTCTTTCGCCTGTCTCTGGCAG |
| AK308 | Reverse primer to clone GlnZ isoforms into pBR | GAATTCAAAAAAGCCCATGCAGAGATGG |
| AK379 | Forward primer to clone $GlnZ_{194}$-M1 into pBR | GACGTCTTTTACTTCGGCTCCAAACTTTTCGCCTGTCTCTGG |
| AK407 | Forward primer to make a chromosomal deletion of the *glnA* promoter | ACTTTAACTCTCCTGGATTGGTCATGGTCGTCGTGGTAACGAAATCTGCAGTGTAGGCTGGAGCTGCTTC |
| AK415 | Reverse primer to make a chromosomal deletion of the *glnA* promoter | TTACGCAATTTTTCGATCACAACTTTGCCTCAGGCATTAGAAATAGCGCGATGGGAATTAGCCATGGTCC |
| AK318 | Forward primer to delete *glnZ* from the MG1655 chromosome | TGCGTATGACTCCGCATCCGGTAGAGTTTGAGCTGTACTACAGCGTCTAATGACTCTGCTAATACTGTTACGAGTGGCTGCTTACAGCGGTACTCTGCTACCGCTTTTTTGTGTAGGCTGGAGCTGCTTC |
| AK320 | Reverse primer to delete *glnZ* from the MG1655 chromosome | AACGTGAAGTTTGTCAGCTATCTGTAGCCCATCTCTGCATGGGCTTTTTTGTGTAGGCTGGAGCTGCTTC |
| LW069 | Forward primer to clone GlnA promoter into PM1205 | CGAAGCGGCATGCATTTACGTTGACACCATCGAATGGCGCCCATTGAAGCACTATATTGGTGC |
| LW070 | Reverse primer to clone GlnA promoter into PM1205 | TAACGCCAGGGTTTTCCCAGTCACGACGTTGTAAAACGACAGCGGACATACTTTAACTCTCC |
| LW025 | Forward primer to clone GlnZ promoter into PM1205 | TAACGCCAGGGTTTTCCCAGTCACGACGTTGTAAAACGACCATGAATTCTGTTTCCTGTGTGAAATTGTTATCCGCTCACAATTAGCTCAAACTCTACCGGATGCG |
| LW068 | Reverse primer to clone GlnZ promoter into PM1205 | CGAAGCGGCATGCATTTACGTTGACACCATCGAATGGCGCGACCTGCCGCCAGAAGAAGC |
| LW189 | Forward primer to replace SucA with *cat-sacB* in the chromosome | AGCGCAGCGCATCAGGCGTAACAAAGAAATGCAGGAAATCAATGAGACGTTGATCGGCACGTAAG |
| LW202 | Reverse primer to replace SucA with *cat-sacB* in the chromosome | GGGACCAGAATATCTACGCTACTCATTGTGTATCCTTTATGTAACAGATGAACAGCATGTAACACC |
| LW234 | Forward primer to amplify pUC19 for Gibson cloning | ATCCGCTTACAGACAAGC |
| LW235 | Reverse primer to amplify pUC19 for Gibson cloning | AGCTGTTTCCTGTGTGAAATTG |
| LW236 | Forward primer to clone SucA-SPA into pUC19 | ATTTCACACAGGAAACAGCTAGCGCAGCGCATCAGGCGTAACAAAGAAATGC |
| LW237 | Reverse primer to clone SucA-SPA into pUC19 | CAGCTTGTCTGTAAGCGGATGGGACCAGAATATCTACGCTACTCATTGTGTATC |
| LW240 | Forward QuikChange primer to make compensatory mutant in pUC19-SucA-SPA | TTTTATGCTTACTTCGCGGTGGATACTACCACGCA |
| LW241 | Reverse QuikChange primer to make compensatory mutant in pUC19-SucA-SPA | TGCGTGGTAGTATCCACCGCGAAGTAAGCATAAAA |
| LW242 | Forward primer to make SucA-SPA/SucA-M1-SPA on the chromosome | AGCGCAGCGCATCAGGCGTAAC |
| LW243 | Reverse primer to make SucA-SPA/SucA-M1-SPA on the chromosome | GGGACCAGAATATCTACGCTACTCATTGTG |

| | | |
|---|---|---|
| LW117 | Forward primer to make *glnP*-mCherry on the chromosome of NRD1166 | ATACTATGCCGATATACTATGCCGATGATTAATTGTCAACGCGCTTCAGCCATACTTTTCATACTC |
| LW118 | Reverse primer to make *glnP*-mCherry on the chromosome of NRD1166 | CCTTGATGATGGCCATGTTATCCTCCTCGCCCTTGCTCACAAACTGCATATGTTGTTCCTGTTACCG |
| LW137 | Reverse primer to make *glnP*-M1-mCherry on the chromosome of NRD1166 | CCTTGATGATGGCCATGTTATCCTCCTCGCCCTTGCTCACAAACTGCATATGTTGTTCCTGTTA |

**Table S4.** Summary of GlnZ targets based on RIL-seq experiments. Summary of sRNAs targets based on RIL-seq experiments. RIL-seq datasets from experiments done in six different conditions (11,12) were analyzed in order to predict GlnZ targets. Table is sorted according to the number of conditions in which a target was found. Red font indicates GlnZ was the first RNA in the chimera, for all others GlnZ was the second RNA. Targets also found in the top 50 candidates from the MAPS dataset are highlighted in grey.

**Table S5.** Summary of GlnZ targets based on MAPS experiments. MS2 affinity purification followed by RNA-seq was performed with stationary phase cultures grown in Gutnick medium with 0.4% glucose and 15 mM ammonium in duplicate. Differential expression analysis was carried out using DESeq2. Table is sorted by most significant adjusted p-value (padj). lfcSE corresponds to the standard error of the $log_2$-fold change. Targets also found in the top 50 of the RIL-seq dataset are highlighted in grey.
